# Sterilizing immunity against SARS-CoV-2 in hamsters conferred by a novel recombinant subunit vaccine

**DOI:** 10.1101/2020.12.18.423552

**Authors:** Yangtao Wu, Xiaofen Huang, Lunzhi Yuan, Shaojuan Wang, Yali Zhang, Hualong Xiong, Rirong Chen, Jian Ma, Ruoyao Qi, Meifeng Nie, Jingjing Xu, Zhigang Zhang, Liqiang Chen, Min Wei, Ming Zhou, Minping Cai, Yang Shi, Liang Zhang, Huan Yu, Junping Hong, Zikang Wang, Yunda Hong, Mingxi Yue, Zonglin Li, Dabing Chen, Qingbing Zheng, Shaowei Li, Yixin Chen, Tong Cheng, Jun Zhang, Tianying Zhang, Huachen Zhu, Qinjian Zhao, Quan Yuan, Yi Guan, Ningshao Xia

## Abstract

A safe and effective SARS-CoV-2 vaccine is essential to avert the on-going COVID-19 pandemic. Here, we developed a subunit vaccine, which is comprised of CHO-expressed spike ectodomain protein (StriFK) and nitrogen bisphosphonates-modified zinc-aluminum hybrid adjuvant (FH002C). This vaccine candidate rapidly elicited the robust humoral response, Th1/Th2 balanced helper CD4 T cell and CD8 T cell immune response in animal models. In mice, hamsters, and non-human primates, 2-shot and 3-shot immunization of StriFK-FH002C generated 28- to 38-fold and 47- to 269-fold higher neutralizing antibody titers than the human COVID-19 convalescent plasmas, respectively. More importantly, the StriFK-FH002C immunization conferred sterilizing immunity to prevent SARS-CoV-2 infection and transmission, which also protected animals from virus-induced weight loss, COVID-19-like symptoms, and pneumonia in hamsters. Vaccine-induced neutralizing and cell-based receptor-blocking antibody titers correlated well with protective efficacy in hamsters, suggesting vaccine-elicited protection is immune-associated. The StriFK-FH002C provided a promising SARS-CoV-2 vaccine candidate for further clinical evaluation.

## Introduction

The first coronavirus pandemic in human history, caused by severe acute respiratory syndrome coronavirus 2 (SARS-CoV-2), is changing the whole landscape of global public health. To date, 218 countries and regions around the world have confirmed SARS-CoV-2 infections, with over 60 million confirmed COVID-19 cases and costing nearly 1.5 million people’s lives, and the number of human infections is still rocketing up. Even though many efforts could slow down the virus transmission, it is generally believed that the ultimate tool to avert such a pandemic should be an effective and safe vaccine, with which to achieve herds immunity. Thus, the advent of one or more efficacious vaccines, having the availability, accessibility, and affordability across the globe, is essential for achieving effective control of the pandemic, which is still on-going even with new peaks of all-time high of 2^nd^ wave in the winter in certain countries/areas of the northern hemisphere.

The cellular entry of SARS-CoV-2 is predominantly mediated by the interaction of viral spike protein and its major receptor, the host angiotensin-converting enzyme 2 (ACE2) (*1-3*). Like other coronaviruses, the SARS-CoV-2 spike is a homotrimer protruding from the viral surface, and it comprises two functional subunits, namely, the S1 subunit containing the receptor-binding domain (RBD) and the S2 subunit mediating membrane fusion between the virus and the cell (*4-6*). To date, several lines of evidence showed that antibodies against this spike protein could play a critical role in the immunoprophylaxis and therapeutics against COVID-19 (*7-12*). A vast array of different platforms with different molecular modality is currently being employed for vaccine development against COVID-19 (*13-19*). Subunit vaccines, particularly in the case that protein making up the subunits could be prepared using recombinant DNA technology, are of great advantage due to its proven safety and compatibility with multiple boosts if necessary (*20*).

Two challenges, however, should be addressed to develop an ideal SARS-CoV-2 subunit vaccine. The first is to prepare the recombinant protein subunits in a functional form that can predominantly elicit SARS-CoV-2 neutralizing antibodies rather than non-neutralizing antibodies, as the latter may mediate antibody-dependent enhancement (ADE) for disease or infection (*21*). The second is to overcome the less immunogenic drawback of the subunit protein than an intact virus or virus-like particle-based vaccines (*20*). As the classical Alum adjuvant is generally apt to stimulate Th2-biased immune response, its use could be associated with vaccine-associated enhanced respiratory disease (VAERD) (*22, 23*). New adjuvants that can help to generate a more balanced Th1/Th2 cellular response and to improve the immunogenicity of the non-particulate protein-based vaccine are certainly required.

In this study, through *in vivo* evaluation of the immunogenicity of CHO-expressed various spike derivatives, we identified the full length of the spike ectodomain (StriFK), which induced more neutralizing to non-neutralizing antibodies, as the best performing immunogen. In mice, hamsters, and non-human primates, we tested the immunostimulatory effects of different adjuvants with StriFK, specifically FH002C (an nitrogen bisphosphonate-modified zinc-aluminum hybrid adjuvant, developed in house) and Al001 (a traditional Alum adjuvant commonly used in human vaccines). Enhanced immunogenicity of StriFK was observed in combined use with FH002C compared to that with Al001 for both the functional antibodies and cellular responses. More importantly, we demonstrated the immunizations of StriFK-FH002C in hamsters provided adequate and immune-correlated protection against the infection, pathogenesis, and inter-animal transmission of SARS-CoV-2. Together, our results showed robust immune responses and protective potency elicited by a new adjuvant formulated StriFK vaccine in animals.

## Results

### Comparisons of the antibody response induced by various vaccine candidates in mice

Using the CHO expression system, recombinant spike protein derivatives of SARS-CoV-2, including RBD, S1, S2, trimerized RBD (RBDTfd) and S-ectodomain (StriFK), were obtained (Figure 1A). All five purified proteins achieved acceptable purity and reacted well with antibodies in the COVID-19-convalescent human plasma in western blots (Figure 1B). For adjuvant used in the immunogenicity evaluation in mice (Figure 1C), both the traditional Alum (Al001) and the novel FH002C adjuvants showed similar morphology and antigen adsorption capacity (Figure S1).

**Figure 1.**
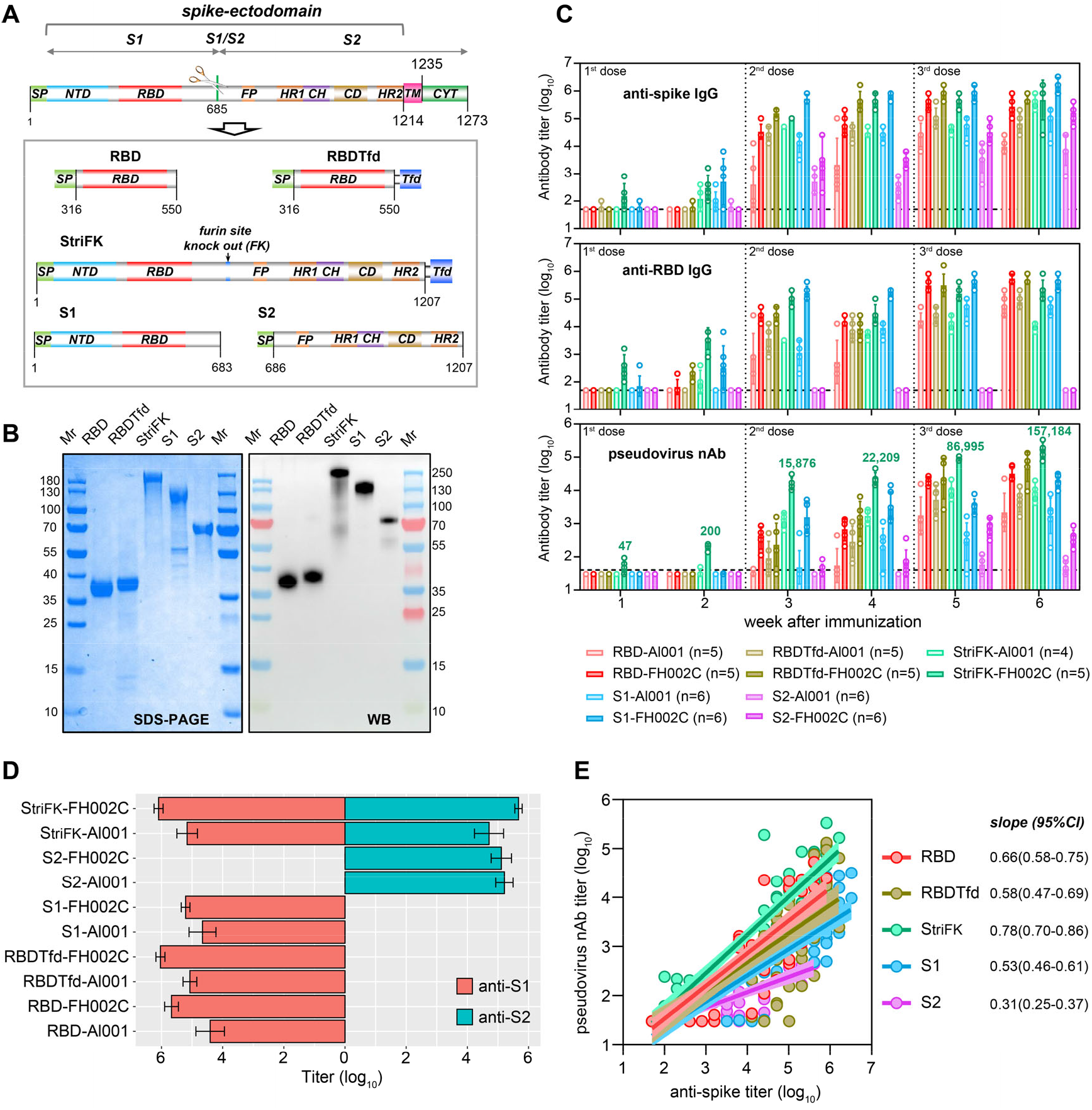
Immunogenicity of various recombinant protein-based vaccine candidates in mice. (A) Schematics of constructs for producing different recombinant spike derivatives. Functional domains of the SARS-CoV-2 spike are colored. NTD, N-terminal domain; RBD, receptor-binding domain; FP, fusion peptide; HR1/2, heptad repeat 1/2; CH, central helix; TM, transmembrane domain; CYT, cytoplasmic tail; Tfd, trimerization motif of T4 fibritin (foldon). (B) SDS-PAGE and western blot (WB) analyses for purified proteins from CHO cells. RBDTfd, RBD with C-terminal fused Tfd; StriFK, furin site removed spike ectodomain with C-terminal fused Tfd. A convalescent serum sample from a COVID-19 patient was used for WB assay. (C) Antibody response induced by various vaccine candidates in mice. Groups of BALB/c mice were immunized with different vaccine candidates (1 μg/dose) at weeks 0, 2, and 4. The proteins of RBD, RBDTfd, StriFK, S1 and S2 were used as the immunogen combined with traditional (Al001) or modified (FH002C) Alum adjuvants, respectively. Mouse serum titers of anti-spike (upper panel), anti-RBD (middle panel), and SARS-CoV-2 pseudovirus neutralizing antibody (lower panel) were displayed. (D) The serum titers of anti-S1 and anti-S2 in immunized mice at Week 6 since the initial immunization. (E) The associations between the titers of pseudovirus nAb and anti-spike binding antibody in mice received different immunogens. The linear regression curves with 95% confidence intervals were plotted. The slopes of regression curves were indicated to reflect the relative ratio of neutralizing-to-binding antibody induced by various immunogens. For panel (C) and (D), data were plotted as the geometric mean±SD.

Among all tested candidates, the StriFK-FH002C immunized mice presented the most rapid antibody response, showing detectable anti-spike, anti-RBD, and neutralization antibodies (nAb) just 1-week after administration of the 1^st^ dose (Figure 1C). Two weeks after the 1^st^ dose, mice receiving StriFK or S1 (with either adjuvant) all presented detectable (geometric mean titer, GMT≥100) anti-spike antibodies, whereas mice received RBDTfd-FH002C, StriFK-Al001, StriFK-FH002C, and S1-FH002C showed detectable anti-RBD antibodies. However, at this time point, only mice immunized with StriFK-FH002C presented pseudovirus neutralizing antibody (GMT=200). Compared to Al001, the new FH002C adjuvant stimulated significantly higher and more rapid antibody response in mice, regardless of the immunogen used. At Week 6, after 3-dose immunization was completed, the RBD, RBDTfd, S1, and StriFK induced similarly high levels of anti-spike and anti-RBD in combination with FH002C adjuvant.

As expected, anti-S2 antibodies were only observed at comparable levels in mice immunized with StriFK and S2 (Figure 1D). Sera from S2-immunized mice showed detectable nAb levels, suggesting that both S1- and S2-specific antibodies contribute to the neutralization activities of StriFK-induced antibodies. Regression analyses on the associations between the anti-spike binding titer and the neutralizing titer in mice receiving various immunogens revealed that the StriFK had a higher neutralizing-to-binding ratio (as reflected by a larger slope value) than the others (Figure 1E).

In a stable pool of CHO cells transfected with StriFK-expressing construct, protein expression reached over 200 mg/L (Figure S2A). The molecular weight of the StriFK was determined to be approximately 700-kDa by HPLC-SEC analysis (Figure S2B). The StriFK showed a binding affinity of 3.52 nM to human ACE2 protein (Figure S2C) and exhibited good reactivities to human COVID-19-convalescent plasmas in ELISA-binding assay (Figure S2D). The StriFK-binding activities of human COVID-19-convalescent plasmas positively correlated with their neutralizing titers against pseudotyped and authentic SARS-CoV-2 virus, also well associated with their receptor-blocking titers (Figure S2E).

Together, the StriFK combined with the FH002C adjuvant provided an excellent vaccine candidate for further evaluation.

### StriFK-FH002C stimulated potent SARS-CoV-2 specific T cell response in mice

To test the vaccination effect in promoting the germinal center formation, we measured the frequencies of T follicular helper (Tfh), germinal center B (GCB), and plasma cells in lymph nodes (LNs) in mice (1 week after a single immunization). In comparison to StriFK-Al001, the StriFK-FH002C significantly increased the frequencies of Tfh (Figure 2A), GCB (Figure 2B), and plasma cells (Figure 2C), which were consistent with the more rapid antibody response and earlier antibody affinity maturation (indicated by higher IgG avidity) after StriFK-FH002C immunization (Figure S3A).

**Figure 2.**
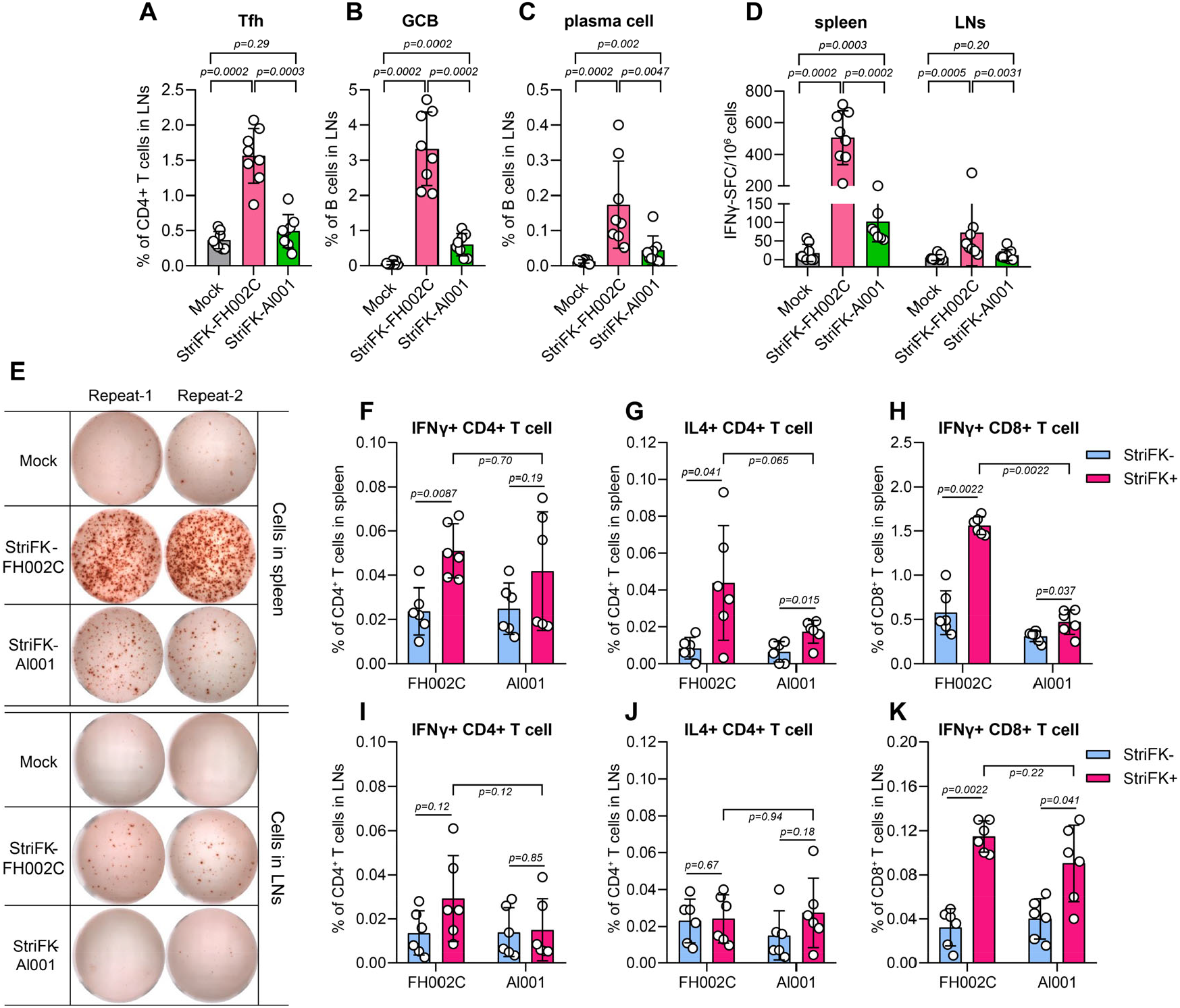
The StriFK-FH002C vaccine candidate elicited a potent humoral and cellular immune response against the SARS-CoV-2 spike in mice. The frequencies of follicular helper T (Tfh) cell (A), germinal center B (GCB) cell (B), and plasma cell (C) that induced by StriFK-FH002C and StriFK-Al001 immunizations. C57BL/6 mice (n=8/group) were immunized with StriFK-FH002C and StriFK-Al001 (both at 10 μg/dose), and lymphocytes in lymph nodes (LNs) were collected at day 7 after the immunization for FACS. (D) and (E) IFN-γ ELISPOT assays for lymphocytes in spleen and LNs under pre-stimulation of a peptide pool covering the entire spike protein. (D) Quantitative measurements of the numbers of IFN-secreting cells. (E) Representative images of ELISPOT detections. (F) Intracellular cytokine staining assays for IFN-γ+CD4+, IL4-γ+CD4+, and IFN-γ+CD8+ T cells of the splenocytes (F, G, and H) or lymphocytes of LNs (I, J, and K) in response to spike peptide pool. The data were plotted as the mean±SD. The Mann-Whitney U test was used for inter-group statistical comparison.

In mice receiving 2-dose immunizations of StriFK-FH002C and StriFK-Al001, we further measured spike-specific T cell response. In contrast to non-immunized animals, the IFNγ ELISPOT assays revealed the numbers of IFNγ-secreting cells after *ex vivo* antigen-peptides stimulation increased by 28.9-fold and 5.8-fold in the spleen, and 14.0-fold and 2.3-fold in lymph nodes (LNs), in mice immunized by StriFK-FH002C and StriFK-Al001, respectively (Figure 2D and 2E). Compared to Al001, the FH002C adjuvant elicited significantly higher frequencies of spike-specific IFNγ-secreting cells in either spleen (p=0.0002) or LNs (p=0.0031).

Intracellular cytokine staining (ICS) measurements of mouse splenocytes stimulated by spike-peptides demonstrated that the StriFK-FH002C successfully induced Th1, Th2, and CTL responses, which were evidenced by significantly higher frequencies of spike-specific T cells of IFN-γ^+^CD4^+^ (p=0.0087, Figure 2F), IL4^+^CD4^+^ (p=0.041, Figure 2G) and IFN-γ^+^CD8^+^(p=0.0022, Figure 2H), respectively. The StriFK-Al001 also increased spike-specific IL4^+^CD4^+^ (p=0.015, Figure 2G) and IFN-γ^+^CD8^+^ T cells (p=0.037, Figure 2H) in splenocytes, but it appeared less potent than StriFK-FH002C. In spike-peptide re-stimulated cells from LNs, although the increases of T helper cells (IFN-γ^+^CD4^+^ and IL4^+^CD4^+^) were not pronounced (Figure 2I and 2J), markedly elevated IFN-γ^+^CD8^+^ T cells were observed in both vaccinated groups (Figure 2K).

Consistent with T cell analysis findings, sera from StriFK-FH002C immunized animals showed significantly higher IgG2a or 2b titers (Figure S3B) and IgG2-to-IgG1 titer ratio (Figure S3C) than StriFK-Al001. Together, these results demonstrated the StriFK-FH002C could induce a more balanced Th1/Th2 immune response.

### Immunogenicity of StriFK-FH002C in non-human primates and hamsters

Next, we examined the immunogenicity of StriFK-FH002C in cynomolgus macaques in comparison to StriFK-Al001 and RBD-FH002C (Figure S4). After the 2-dose regimen, at Week 4 all vaccine-immunized monkeys showed detectable anti-spike (Figure S4A), neutralizing (Figure S4B), and receptor-blocking antibodies (Figure S4C). In contrast to RBD, the StriFK stimulated higher antibody titers in all three assays, particularly for the 1^st^ and 2^nd^ dose. On the other hand, the FH002C adjuvant showed advantages in accelerating antibody generation. After the 3-dose immunization completion, animals immunized with all three vaccine candidates displayed similarly high levels of nAb and receptor-blocking antibodies, suggesting the advantages of StriFK-FH002C were mainly reflected in the rapid antibody response in the early immunization phase. However, the highest neutralizing response (GMT=32,081, one was 26,721, and the other one was 38,515) was still observed in monkeys received 3-dose of StriFK-FH002C immunization at Week 7. No abnormality in body weight (Figure S4D), body temperature (Figure S4D), liver function (Figure S4E), renal function (Figure S4F), cardiac enzymes (Figure S4G) or other clinical manifestation was observed in StriFK-FH002C or StriFK-Al001 immunized animals, suggesting a good safety profile of these vaccines in non-human primates.

As non-human primate was reported as a mild disease model for SARS-CoV-2 infection (*24*), while hamster is considered as a suitable animal model mimicking severe COVID-19 pneumonia in humans (*25-27*), here we assessed the immunogenicity of vaccine candidates in Syrian golden hamsters (*Mesocricetus auratus*), aiming to evaluate the protective efficacy using this model (Figure 3). Hamsters received StriFK-based vaccines (either StriFK-FH002C or StriFK-Al001), quickly generated detectable antibodies as early as Week 1 after 1^st^-shot in all four antibodies measured. At Week 3, 7 days after the 2^nd^ dose regimen, the StriFK-FH002C elicited the highest titers of anti-RBD (GMT=462,178; range: 102,400-1,968,300), anti-spike (GMT=1,604,827; range: 409,600-17,714,700), neutralizing antibody (GMT=16,617; range: 6,478-31,220), and receptor-blocking antibody (GMT=634; range: 369-1,537). From Weeks 3 to 6, the antibody titers in StriFK-FH002C or StriFK-Al001 immunized animals slightly decreased to about 50% of the corresponding peak values. When a booster immunization was given at Week 6, the serum antibodies of animals in groups of StriFK-FH002C and StriFK-Al001 rapidly increased to comparable levels as that at Week 3. In hamsters, the StriFK-FH002C induced more than 4-fold higher neutralizing receptor-blocking antibody titers than StriFK-Al001, implying the potent immunostimulatory effect of the FH002C adjuvant. Surprisingly, the RBD-FH002C, which could generate binding and functional antibodies in mice and monkeys, showed very little even no antibody response in hamsters.

**Figure 3.**
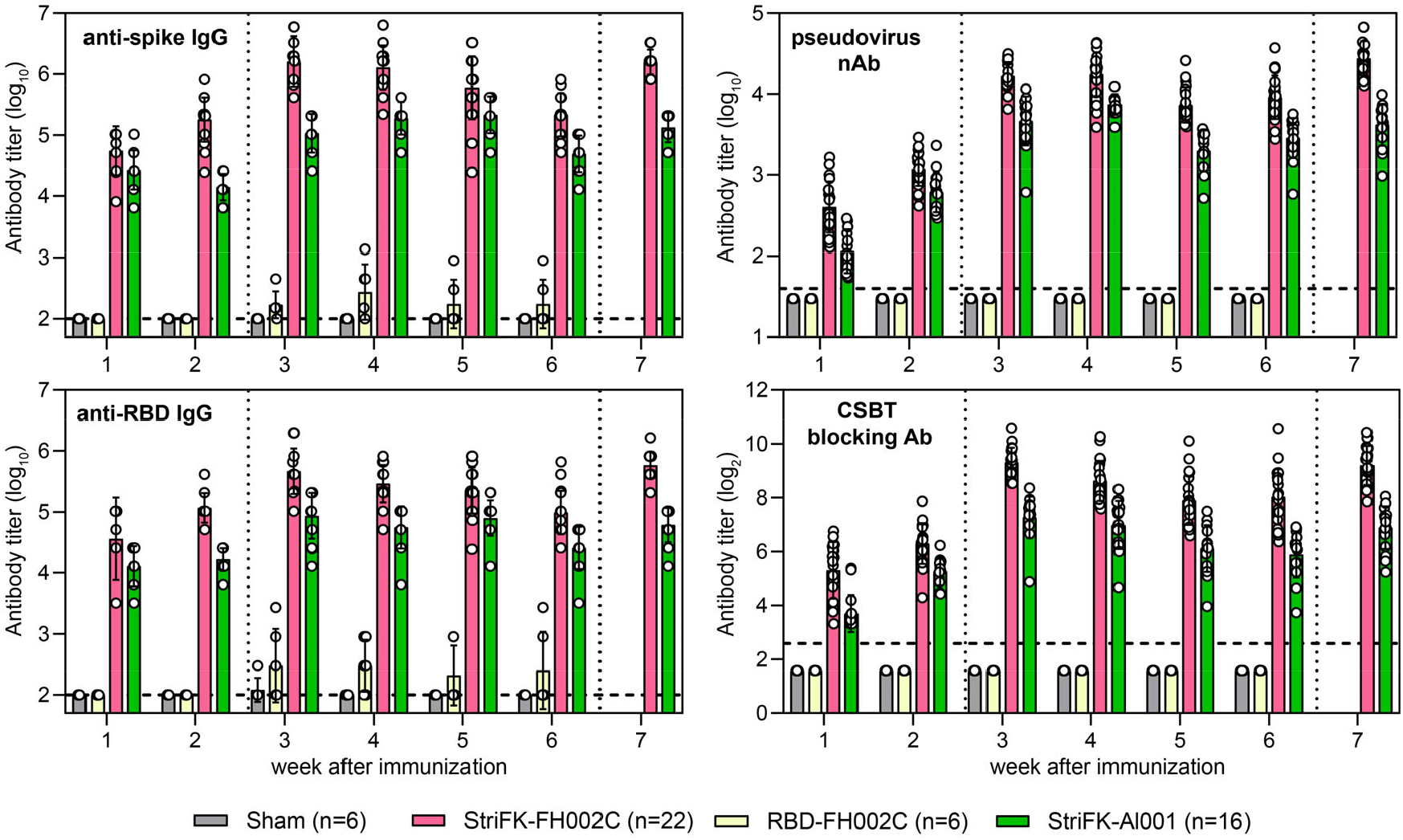
Antibody response induced by immunizations of StriFK-FH002C, RBD-FH002C, and StriFK-Al001 vaccine candidates in hamsters. All three vaccines were used at 10 μg/dose. Animals in Sham (FH002C-only) and RBD-FH002C groups received 2-shot immunizations at Weeks 0 and 2, whereas animals in the StriFK-Al001 group received 3-shot immunizations at Weeks 0, 2, and 6. In the StriFK-FH002C group, 6 of 22 animals received 2-shot immunizations at Weeks 0 and 2, and the remaining 16 animals were immunized for 3-dose (10 μg/dose) at Weeks 0, 2, and 6. Serum samples were collected every week since initial immunization, and the anti-spike, anti-RBD, neutralizing antibody, and the receptor-blocking antibody titers were measured. The quantitative data of antibody titers were plotted as the geometric mean±SD.

### Protective efficacy of StriFK-FH002C against SARS-CoV-2 infection and pathogenesis in hamsters

Using the intranasal SARS-CoV-2 challenging method, we investigated the protective efficacy of StriFK-FH002C in hamsters. We first tested whether a 2-shot immunization of StriFK-FH002C could protect hamsters from SARS-CoV-2 infection and its related pneumonia. In this experiment, the neutralizing antibody GMT of the six StriFK-FH002C vaccinated hamsters was 14,404 (range: 5,368-37,078). In the sham group that received FH002C-only, hamsters lost an average of 8.4%, 12.2%, and 11.7% of body weight by 3, 5, and 7 days post-infection/inoculation of virus (dpi), respectively (Figure 4A). In contrast, StriFK-FH002C immunized animals lost just an average of 2.0%, 2.1%, and 1.2% of body weight by the corresponding time points (Figure 4A), which was significantly lower than that of the sham controls (p<0.0001, two-side two-way ANOVA test) and similar to that of the uninfected (mock) group (p=0.061). All animals in the sham group exhibited obvious symptoms of weakness, piloerection, hunch back, or abdominal respiration, which was observed in neither vaccinated nor un-infected animals (Figure 4B).

**Figure 4.**
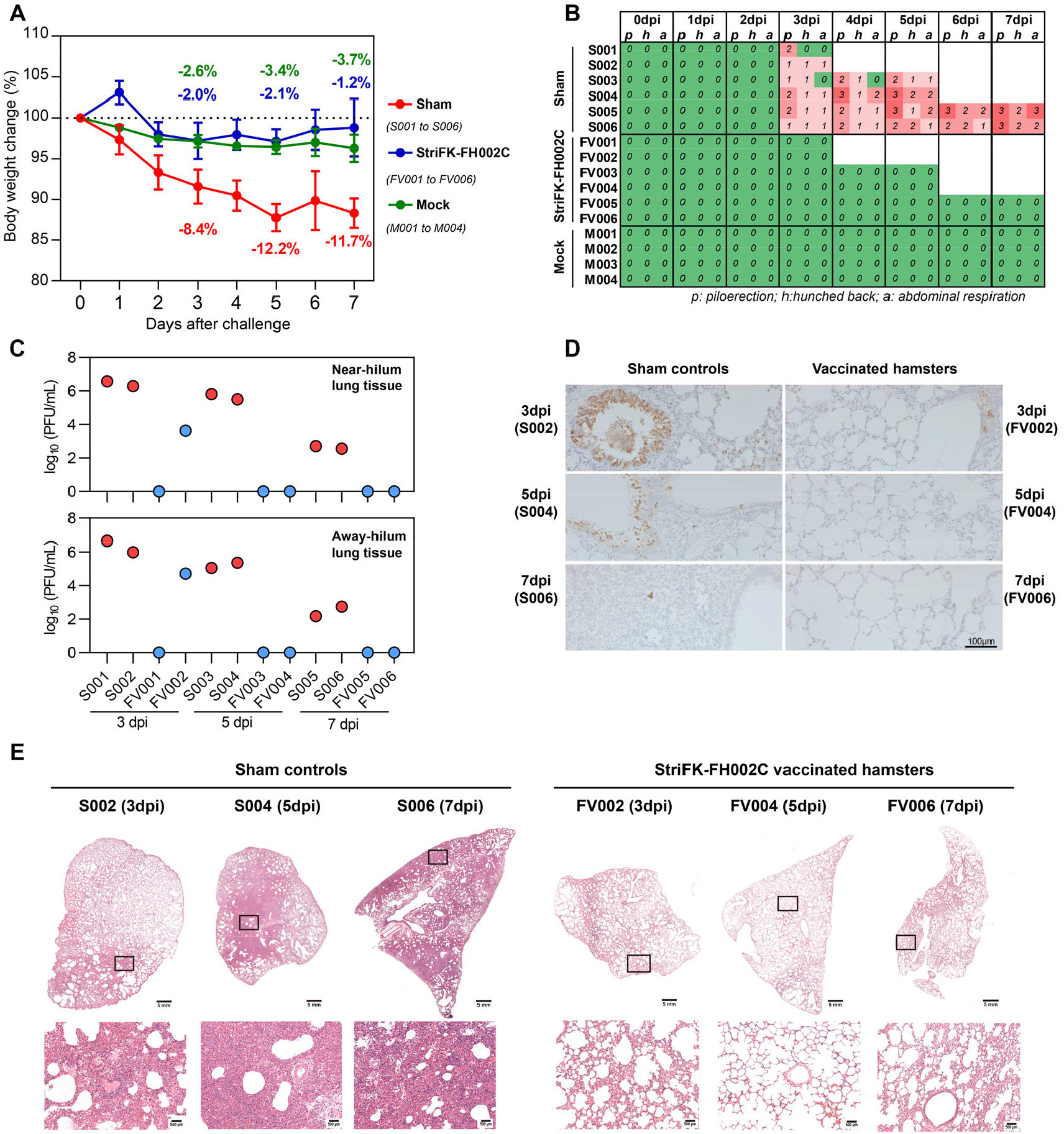
Protective efficacy of 2-shot immunizations of StriFK-FH002C against intranasal SARS-CoV-2 challenge in hamsters. Six hamsters in StriFK-FH002C group (FV001 to FV006) and 6 in Sham group (S001 to S006), were intranasally challenged with 1×10^4^ PFU of SARS-CoV-2. Additional 4 hamsters (M001 to M004) were used as uninfected controls (mock). On days 3, 5, and 7 post virus challenge, two hamsters in StriFK-FH002C and Sham groups were sacrificed for tissue analyses, respectively. (A) Changes in body weight following virus challenge. Data were shown as mean±SD. The average weight loss of each group was indicated as a colored number. (B) Scores of typical symptoms (piloerection, hunched back, and abdominal respiration) of hamsters following virus challenge. The symptoms were scored based on the severity of none (0), moderate (1), mild (2), severe (3) and, critical (4). (C) Infectious virus titers in the lungs were collected from tissues near (upper panel) or away (lower panel) from the pulmonary hilum at 3, 5, and 7 days post-inoculation of the virus (dpi). (D) Viral N protein detected in the lungs from virus-challenged hamsters. (E) H&E staining for lung sections collected from virus-challenged hamsters. Views of the whole lung lobes were presented in the upper panel, and the areas in the black box were enlarged in the lower panel.

The 2-shot immunized animals in sham and vaccine groups were executed at 3, 5, and 7 dpi (2 animals per time point) to assess the viral loads and histopathological profiles in the respiratory tract. For sham controls, the peak viral loads in lung tissues were detected at 3 dpi (6.4±0.3 log_10_ PFU/mL) and gradually decreased at 5 dpi (5.4±0.3 log_10_ PFU/mL) and 7 dpi (2.5±0.3 log_10_ PFU/mL). In the StriFK-FH002C group, the infectious virus was only detected in lung tissues of hamster FV002 (4.2 log_10_ PFU/mL) at 3 dpi, but not in tissues of other animals (Figure 4C). The relatively lower neutralizing antibody titer in FV002 (ID_50_=5,368) than the others (ID_50_ range: 9,363-37,078) was a possible explanation for the insufficient protection of this animal individual. However, the lung viral titer of the FV002 hamster remained markedly lower than those from sham controls. The nucleocapsid immunostainings in the lungs were largely consistent with viral titer profiles (Figure 4D). Pathological examination of the lung sections by hematoxylin and eosin (H&E) staining demonstrated typical features of moderate-to-severe interstitial pneumonia in sham controls. In contrast, vaccinated animals only showed slight histological changes with significantly reduced pathological severity (Figure 4E).

To further evaluate the protective efficacy of StriFK-FH002C and to reveal the role of FH002C adjuvant, 3-shot immunizations of StriFK-FH002C or StriFK-Al001 (n=8 each) were conducted in hamster for the SARS-CoV-2 challenging study (Figure 5A). The neutralizing and receptor-blocking antibody titers in these StriFK-FH002C vaccinated animals were approximately 4 folds higher than that in StriFK-Al001 vaccinated animals. The average weight loss at 5 dpi was 9.0%, −0.04% and 0.17% in the unvaccinated controls, the StriFK-FH002C and the StriFK-Al001 groups, respectively (Figure 5B). The weight loss was markedly lower in both vaccinated groups compared to unvaccinated controls (p<0.0001 for both comparisons) and showed no significant difference between the StriFK-FH002C and StriFK-Al001 groups (p=0.39). Infectious viruses were detected in all unvaccinated animals’ lungs. Interestingly, the males showed higher viral titers than females. In the lung tissues of vaccinated hamsters, no infectious virus was detected (Figure 5C).

**Figure 5.**
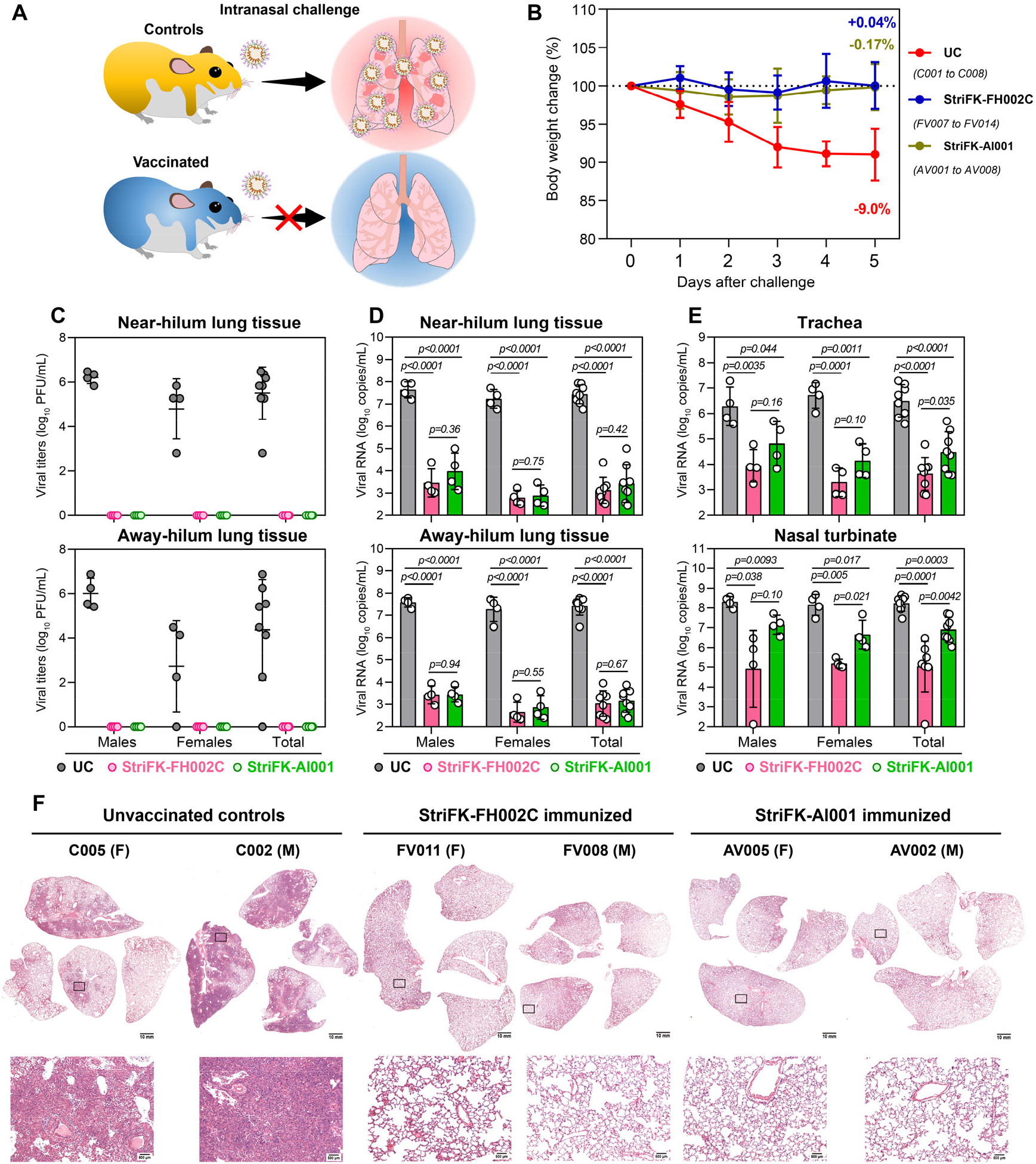
Comparison of the protective efficacy of 3-shot immunizations of StriFK-FH002C and StriFK-Al001 against intranasal SARS-CoV-2 challenge in hamsters. (A) Schematics of the virus challenging study. A total of 24 hamsters (half were males and half were females), including 8 unvaccinated animals (UC, C001 to C008) and 16 immunized animals, which received 3-shot StriFK-FH002C (n=8, FV007 to FV014) or StriFK-Al001 (n=8, AV001 to AV008), were intranasally challenged with 1×10^4^ PFU of SARS-CoV-2. All animals were sacrificed for tissue analyses at 5 dpi. (B) The changes in body weight following virus challenge. The average weight loss of each group at 5 dpi was indicated as a colored number. Data were shown as mean±SD. (C) Infectious viral titers in the lungs were collected from tissues near (upper panel) or away (lower panel) from the pulmonary hilum. (D) Viral RNA levels in the lungs were collected from tissues near (upper panel) or away (lower panel) from the pulmonary hilum. (E) Viral RNA levels in lysates of the trachea (upper panel) and nasal turbinate (lower panel) tissues from hamsters. Data shown were mean±SD. The Mann-Whitney U test was used for inter-group statistical comparison. (D) H&E staining for lung sections collected from virus challenged hamsters. Views of the whole lung lobes (4 independent sections) were presented in the upper panel, and the areas in the black box were enlarged in the lower panel.

Consistent with infectious virus titers, viral RNA levels were profoundly reduced (over 4 logs) in lung tissues from vaccinated hamsters compared to the controls (Figure 5D). In trachea and nasal turbinate tissues, StriFK-FH002C immunized hamsters exhibited more marked viral RNA reductions (over 3 logs) than those received StriFK-Al001 (Figure 5E). No significant gender difference was found for vaccination-mediated viral RNA reduction. Gross lung pictures demonstrated multifocal diffuse hyperemia and consolidation in the lungs of controls, and the apparent lesions were significantly diminished in both vaccinated groups (Figure S5A). Histopathological examinations observed pulmonary consolidation, alveolar destruction, and diffuse inflammation in about 20-30% of lungs from female controls and 50-60% of lungs from male controls, suggesting male hamsters had more severe pneumonia than females. Comparing to controls, vaccinated animals (using StriFK-FH002C or StriFK-Al001), either males or females, only showed mild inflammation in limited areas of lungs, and had minimal pneumonia pathology in most areas of lungs (Figure 5F).

These results demonstrated StriFK-FH002C immunizations effectively prevented hamsters from SARS-CoV-2 virus infection and pathogenesis.

### Protective efficacy of StriFK-FH002C against SARS-CoV-2 inter-animal transmission in hamsters

To mimic the disease transmission, we also assessed whether the StriFK-FH002C vaccine could protect inter-animal SARS-CoV-2 transmission in hamsters (Figure 6A). Co-housed with the infected donors, the unvaccinated control hamsters showed their maximal weight loss on Day 4 (mean: 8.5%) post-contact exposure (Figure 6B). In contrast, the weight loss was diminished in both vaccinated groups (StriFK-Al001 *vs* controls, p=0.0091; StriFK-FH002C *vs* controls, p<0.0001), and it changed to a lesser degree (p=0.017) in the StriFK-FH002C group (mean: −0.22%) than that in StriFK-Al001 group (mean: 3.6%). None of the vaccinated animals presented infectious virus in their lungs at Day 5 post-exposure, whereas all control animals had detectable infectious viruses at varying titers (Figure 6C). More consistent with the weight change findings, viral RNA detections revealed a lower viral load in hamsters of the StriFK-FH002C group than in StriFK-Al001 immunized animals or unvaccinated controls (Figure 6D and 6E). Gross lung images (Figure S5B) and pathological analyses (Figure 6F) indicated that control hamsters had extensive lung damage, with consolidated lesion and inflammatory cell infiltration across larger areas, which was quite similar to that presented in intranasal SARS-CoV-2 challenged hamsters. In contrast, both vaccines prevented tissue damage to a large degree, if any, with mild perivascular and alveolar infiltration observed in very few areas (Figure 6F).

**Figure 6.**
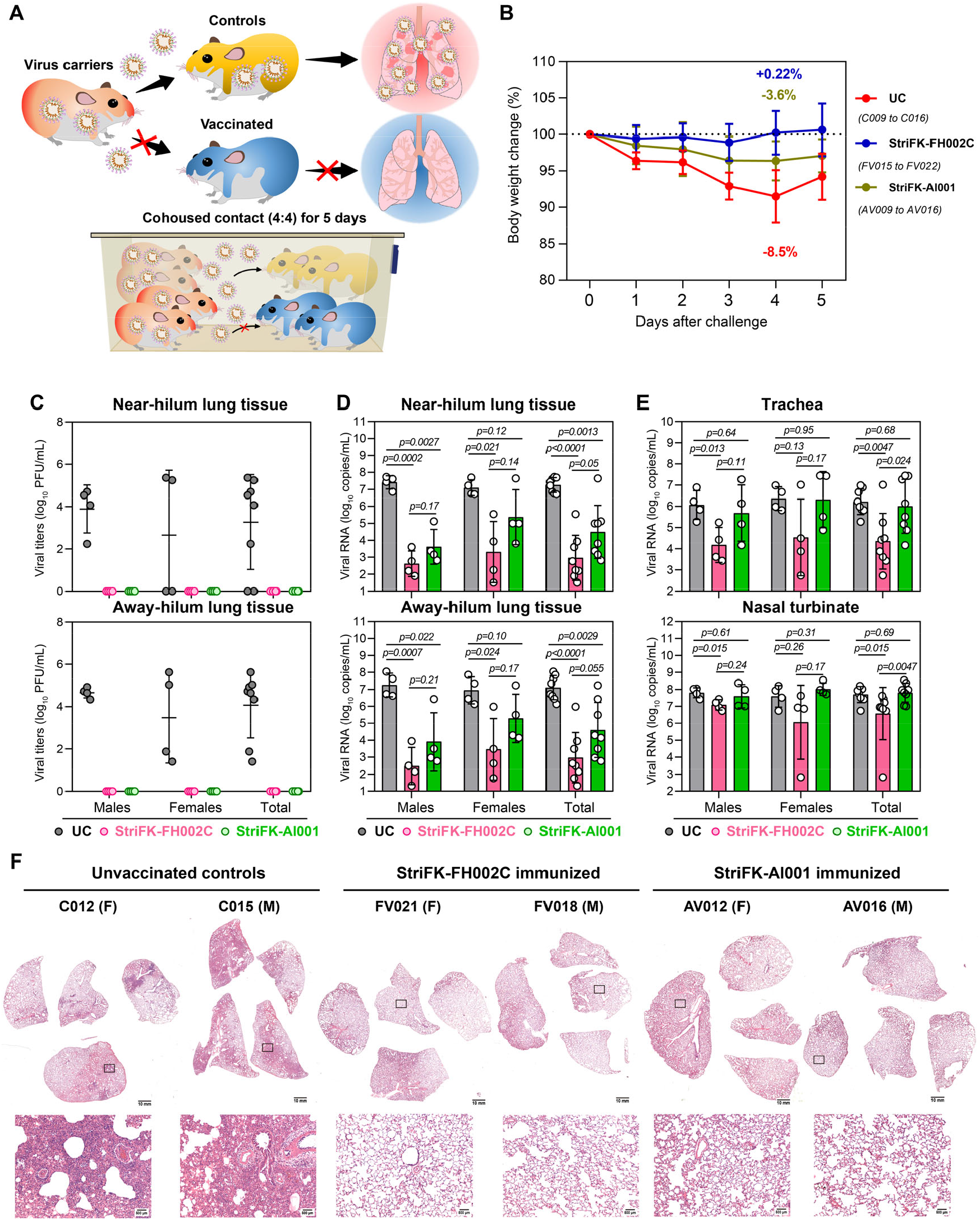
Comparison of the protective efficacy of 3-shot immunizations of StriFK-FH002C and StriFK-Al001 in preventing SARS-CoV-2 transmission in golden hamsters by direct contact. (A) Schematics of virus transmission study. Firstly, 24 hamsters were pre-infected with 1×10^4^ PFU of SARS-CoV-2 as virus donors. One-day later, every 4 donors were co-housed with 4 vaccinated or control hamsters for a 5-day follow-up. A total of 24 hamsters (male:female=1:1), including 8 unvaccinated animals (UC, C009 to C016) and 16 immunized animals, which received either 3-shot StriFK-FH002C (n=8, FV015 to FV022) or StriFK-Al001 (n=8, AV009 to AV016) were exposed to the SARS-CoV-2 infected donors. (B) Changes in body weight following co-housed exposure. The average weight loss of each group at 4 dpi (the time point that controls showed maximum percent weight loss) was indicated as a colored number. Data shown were mean±SD. (C) Infectious viral titers in the lungs were collected from tissues near (upper panel) or away (lower panel) from the pulmonary hilum. (D) Viral RNA levels in the lungs were collected from tissues near (upper panel) or away (lower panel) from the pulmonary hilum. (E) Viral RNA levels in lysates of the trachea (upper panel) and nasal turbinate (lower panel) tissues from hamsters. Data shown were mean±SD. The Mann-Whitney U test was used for inter-group statistical comparison. (D) H&E staining for lung sections collected from tested hamsters on day 5 after co-housed exposure. Views of the whole lung lobes (4 independent sections) were presented in the upper panel, and the areas in the black box were enlarged in the lower panel.

### Immune-correlation of protective efficacy of StriFK-FH002C vaccination

Using pooled animals related to Figures 5 and 6, we analyzed the associations between the StriFK-elicited functional antibody titers and indices for protective efficacy. We demonstrated that the serum titers (before the virus challenge) of neutralizing antibody and receptor-blocking antibody were negatively correlated with viral levels in tissues of nasal turbinate, trachea, and lung from virus-challenged hamsters (Figure S6). We also compared the serum antibody titers of hamsters before and after the SARS-CoV-2 challenge (Figure 7A and 7B). Control hamsters showed an average of 24-fold (p<0.0001) and 5.4-fold (p<0.0001) increase in neutralizing antibody and receptor-blocking antibody, whereas the StriFK-Al001 immunized animals presented only 2.8-fold (p<0.0001) and 1.4-fold (p=0.0066) elevation in corresponding antibody markers. Notably, the StriFK-FH002C immunized hamsters showed no significant antibody increase after the SARS-CoV-2 challenge in either neutralizing antibody (p=0.19) or receptor-blocking antibody (p=0.98), suggesting that sterilizing immunity to SARS-CoV-2 may have been conferred due to immunization in these animals. The pooled antibody data in mice, hamsters, and non-human primates showed that the neutralizing antibodies induced by StriFK-FH002C in animals were about 28- to 269-fold higher than that of the human COVID-19 convalescent plasmas (Figure 7C).

**Figure 7.**
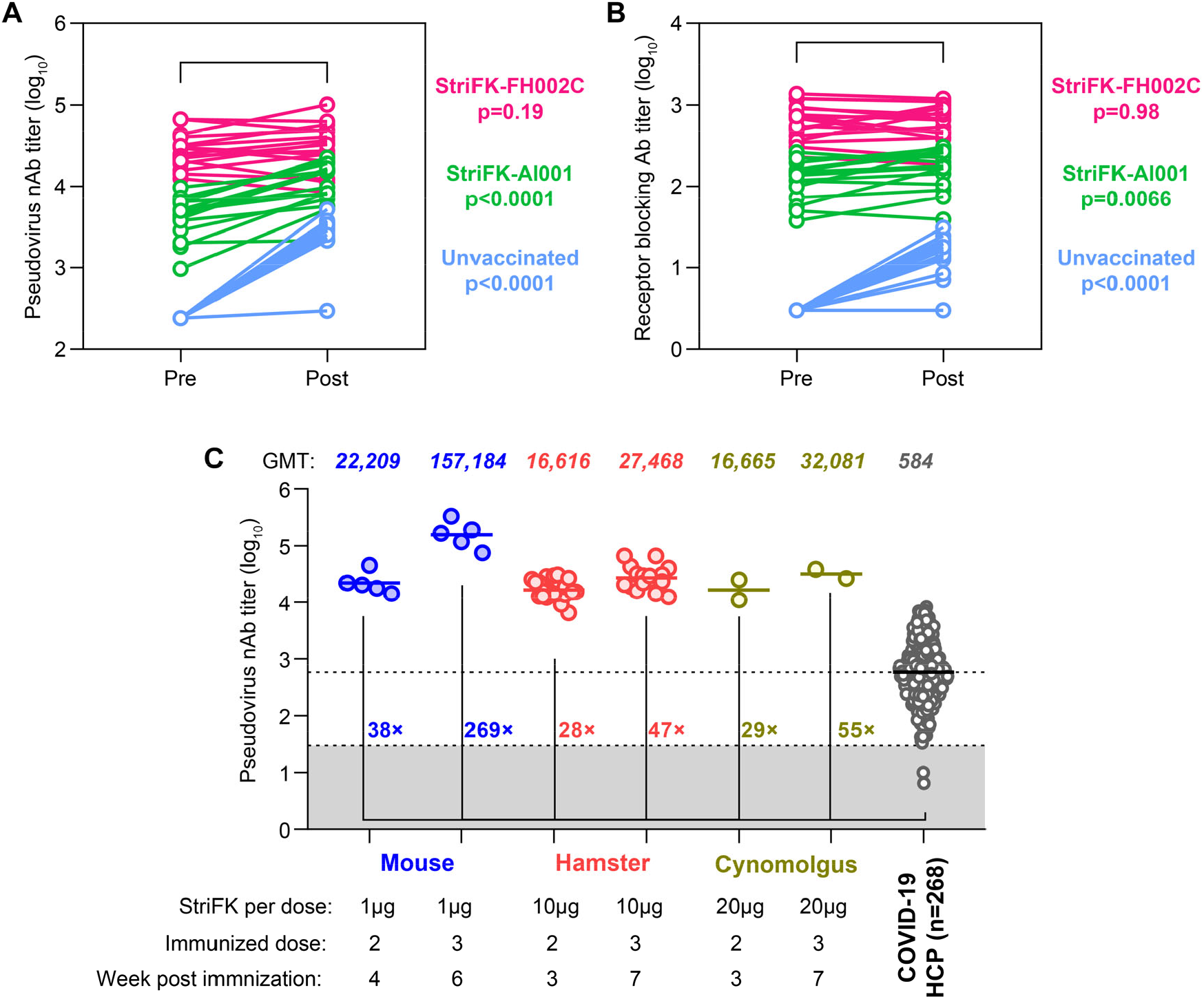
Vaccination with StriFK-FH002C elicited robust protective antibody titers and conferred sterilizing immunity against SARS-CoV-2 in animals. Comparison of the pseudovirus nAb (A) or receptor-blocking antibody (B) titers in paired sera of hamsters collected before (Pre) and 5 days (Post) after SARS-CoV-2 challenges. All vaccinated hamsters and unvaccinated controls involved in experiments related to Figures 5 and 6, were included and pooled for analysis. Paired *t*-test was used for statistical analysis. (C) Comparison of the pseudovirus neutralizing antibody titers in human COVID-19 convalescent plasmas to StriFK-FH002C vaccinated animals. A total of 268 convalescent plasmas collected from recovered COVID-19 patients within 8 weeks since illness onset. The geometric mean titer (GMT) level of pseudovirus nAb of these human samples were 584. The GMT levels of pseudovirus nAb (using the same assay) of StriFK-FH002C vaccinated mice (related to Figure 1, n=6), hamsters (related to Figure 3, n=22) and cynomolgus macaques (related to Figure S3, n=2) were included for comparison.

## Discussion

In this study, we systematically evaluated the immunogenicity and protective efficacy of a recombinant SARS-CoV-2 subunit vaccine candidate SFTK-FH002C in different animal models. This vaccine candidate was composed of CHO-expressed SARS-CoV-2 spike ectodomain protein (StriFK) and a novel adjuvant (FH002C). Comparing to other tested spike derivatives, including RBD, S1, and S2, our *in vivo* data demonstrated the StriFK generated a higher neutralizing-to-binding ratio (Figure 1E), which implied it induced more neutralizing antibodies than non-neutralizing antibodies. The FH002C is a functionalized zinc-aluminum hybrid adjuvant with a significantly improved immune-enhancing effect. Immunization with StriFK-FH002C at low doses in mice, hamsters, and monkeys rapidly generated anti-spike, neutralizing, and receptor-blocking antibodies since one-dose administration. Compared to the GMT level of human COVID-19 convalescent plasmas, 2-shot StriFK-FH002C immunization induced 28-38 folds higher neutralizing antibody and 3-shot immunization generated 47-269 folds higher neutralizing antibody in mice, hamsters, and non-human primates (Figure 7C).

In hamster, an animal model recapitulating severe human COVID-19 disease, StriFK-FH002C vaccination provided protective immunity to prevent the SARS-CoV-2 infection, pathogenesis, and transmission effectively (Figure 4, 5, and 6). The positive correlations between serum functional antibody (neutralizing or receptor-blocking antibody) titers and SARS-CoV-2 virus challenging protective efficacy (Figure S6) demonstrated immune-associated protection of this vaccine. It has been reported that anamnestic responses following virus challenge, which usually occurred in animals with inadequate neutralizing antibody, was a surrogate to evaluate whether the vaccination provides sterilizing immunity (*17*). In our hamster challenging studies, anamnestic responses in either neutralizing or receptor-blocking antibodies were observed in StriFK-Al001 vaccinated-but not in StriFK-FH002C vaccinated-animals (Figure 7A and 7B) following the SARS-CoV-2 challenge, suggesting 3-shot immunizations of StriFK-FH002C should have afforded sterilizing immunity in hamsters. Other evidence to support this was the undetectable infectious virus and the extremely low viral RNA in the lungs of StriFK-FH002C vaccinated animals (Figure 5D and Figure 6D). Although the protective efficacy of StriFK-FH002C in non-human primates was not tested in this study, it can be expected according to the comparable levels of neutralizing and receptor-blocking antibodies induced by this vaccine in cynomolgus macaques to that in hamsters (Figure 7C).

Apart from the highly immunogenic StriFK protein, the novel FH002C adjuvant also plays an essential role in generating high protective antibody and spike-specific T cell response levels. Due to the multiple mechanisms for effective adjuvants, novel adjuvants have focused on different complex formulation involving multiple components with different functions to form various ‘adjuvant systems’ (*28*). In the FH002C, a drug for osteoporosis nitrogen bisphosphonate (risedronate) was loaded on the composite Alum adjuvant by harnessing the Alum’s highly-phosphophilic nature. Zinc ion, one species available to the cellular environments upon adjuvant dissolution, has an immunostimulatory effect and was known to steer the immune response more towards the Th1 pathway (*29, 30*). Although the precise role of nitrogen bisphosphonate was still unknown, previous studies suggested these compounds could stimulate γδ T cells expansion and act as a dendritic cell modulator due to the upstream accumulation of phosphoantigen isopentyl diphosphate as a result of inhibition of farnesyl diphosphate synthetase (*31-33*). Notably, the FH002C adjuvant showed more potent effects than Al001 to promote the developments of Tfh and GCB cells and generated antibody-producing plasma cells in the early days after immunization (Figure 2A, 2B, and 2C). Due to the rapid differentiation of these cell populations, FH002C adjuvant-based StriFK vaccine generated a faster and stronger protective antibody response, which elicited neutralizing antibodies as early as 1-week after the initial immunization. This advantage enables the vaccine’s potential application for emergency vaccination of close contacts due to high-risk exposure, aiming to quickly control outbreaks among close contacts and in the naïve population.

In addition to neutralizing antibodies, T cells are also believed to be essential to vaccine effectiveness (*34-36*). The spike of SARS-CoV-2 has a subnanomolar affinity to its ACE2 receptor, and the infections may occur at the respiratory epithelium of the upper respiratory tract where the antibody concentration is low. Moreover, the functional antibody may not completely block viruses from invading cells; virus-specific T cells may play a remedial role in eradicating virus-infected cells or inhibiting viral replication and spread via non-cytotoxic effects (*37, 38*). Our *in vivo* data showed the StriFK-FH002C immunization could induce potent spike-specific IFNγ+CD8+ and IFNγ+CD4+ T cells, which would help to establish protective immunity afforded by both arms of the immune response in vaccinated individuals.

Safety is also a crucial concern for COVID-19 vaccine development. Previous studies on RSV and measles vaccine suggested VAERD is linked to Th2-biased immune response (*39-41*). A similar phenomenon was also noted in animal studies regarding SARS-CoV vaccines (*39*). Therefore, an ideal COVID-19 vaccine is expected to elicit a balanced Th1/Th2 immune response. Recombinant protein vaccines adjuvanted by traditional aluminum adjuvants generally induce significantly enhanced Th2-biased immune responses but insufficient Th1 responses, which was also observed in the performance of our StriFK-Al001 vaccine in the mouse model, as indicated by the lower IgG2-to-IgG1 titer ratio (Figure S3). The FH002C adjuvant is a novel tool to overcome this drawback. Compared to traditional Alum adjuvant (Al001) used in our licensed HEV and HPV vaccines, the new FH002C adjuvant potently elicited both virus-specific Th1 and Th2 response, as evidenced by the simultaneously elevated IFNγ+CD4+ and IL4+CD4+ T cells (Figure 2F and 2G) in splenocytes of immunized mice. The ICS and ELISPOT assays further demonstrated the StriFK-FH002C could elicit a more robust spike-specific CTL response than the StriFK-Al001 (Figure 2D, 2H, and 2K), which may result from the enhanced Th1 response. On the other hand, as the hamster is a quite susceptible model of SARS-CoV-2 infection and pathogenesis (*25*), all vaccinated hamsters showed significantly diminished virus infection and disease severity without VAERD evidence in our study (Figure 4, 5, and 6). Although the FH002C has not been tested in humans, the risedronate (a distinct component differs from those in Al001) was clinically used to treat osteoporosis over 20-year (*42*). Each dose introduces a fraction (<1/10) of what one would take in an oral dose (e.g., 5 mg) (*42*). In the study, we did not find any systemic side effects in mice, hamsters, or no-human primates (with 2.5 to 3.0 kg macaque receiving a full human dose) immunized with the FH002C adjuvanted vaccine. These results suggested a good safety profile of StriFK-FH002C in animals and supported further clinical trials in humans.

To date, there were over 180 vaccine candidates based on various platforms are under developing against SARS-CoV-2 (*19*). The StriFK-FH002C differs from other reported subunit vaccine candidates in two aspects: the unique FH002C adjuvant and the non-2P mutated StriFK immunogen. FH002C does not rely on natural products (such as squalene, QS-21, etc.). Instead, it utilizes readily available active pharmaceutical ingredients, making it more scalable and more sustainable during the outbreaks and long-term use. On the antigen front, most reported spike-protein-based vaccines were 2P-mutated, but the S-2P production in mammalian cells is difficult with a low expression level (<10 mg/L) in transient transfection according to previous studies (*2, 43*). In our hand, the StriFK expression could reach a level of 40-60 mg/L in transiently transfected CHO cells, whereas the StriFK stably-transfected cell pools expressed the protein as high as >200 mg/L. Cloned high-production cell line showed a yield over 500 mg/L. More importantly, our data demonstrated its promising immunogenicity and protection in animals. The high yield and facile production of StriFK may enable manufacturing robustness and scalability for further applications. Using the CHO expression platform and the novel adjuvant FH002C, various spike proteins from different disease-causing coronavirus strains could be expressed (and kept at the high producing cell pool stage, like ‘cassettes-on-the-shelf’) ahead of time, enabling facile antigen preparation and vaccine formulation toward rapid vaccine development against emerging strains in the future. It is conceivable that this could be an effective approach for controlling the virus spread with rapid development and deployment of an effective vaccine.

In summary, we developed a novel SARS-CoV-2 subunit vaccine based on recombinant StriFK protein and an innovative functionalized adjuvant with unique immunostimulatory properties. Our evaluations demonstrated the StriFK-FH002C was well tolerated and exhibited good immunogenicity in mice, hamsters, and non-human primates. Vaccinations with the StriFK-FH002C provided effective and immune-correlated protectivity against the infection, pathogenesis, and transmission of SARS-CoV-2 in hamsters. Stopping transmission seems to be achievable based on the animal model here. This vaccine candidate is under development in the pre-clinical stage and could be manufactured on a large scale, with the hope that a stronger and more sustainable immune response could be elicited using this vaccine than the natural infection.

## Materials and Methods

### Constructs for recombinant protein expression

The codon-optimized RBD cDNA of SARS-CoV-2 (GenBank: MN908947.3) was obtained by primer annealing. RBDTfd gene was produced by fusing with a trimeric foldon of the T4 phage head fibritin at C-terminal. SARS-CoV-2 S1 and S2 subunit gene referring to the full-length spike glycoprotein gene of SARS-CoV-2 (GenBank: MN908947.3) were synthesized by General Biosystems Co. Ltd (Anhui, China). Spike glycoprotein ectodomain was introduced with three amino acid mutations in furin-like cleavage site that mutating RRAR to GSAS, and fused with a trimeric foldon at C-terminal. All cDNA above were introduced with a C-terminal polyhistidine sequence and codon-optimization was performed for mammalian cell expression. These target genes were constructed into a modified PiggyBac (PB) transposon vector EIRBsMie using NEBuilder HiFi DNA Assembly Master Mix (New England Biolabs, E2621L) described previously (*44*).

### Protein expression and purification

For initial immunogenicity evaluation in mice, the recombinant proteins of RBD, RBDTfd, S1, S2, and StriFK were transiently expressed by ExpiCHO-S cells. The plasmid expression plasmid carrying each SARS-CoV-2-associated protein was transiently transfected into ExpiCHO-S cells by using the ExpiFectamine™ CHO transfection kit according to the manufacturer’s instruction (Thermo Scientific, A29129). After transfection, the cells were placed in an incubator shaker (Kühner AG, SMX1503C) at 37°C containing 8% CO_2_ for 24 hours, and then, transferred the cells to a shaker at 32°C containing 5% CO_2_ culture for 5-7 days. Subsequently, the culture supernatants of CHO cells were subjected to purification by Ni Sepharose Excel column (Cytiva). For further animal vaccination studies, polyhistidine-free StriFK proteins were produced by a stable cell pool transfected with StriFK-expressing construct. The proteins were purified from culture supernatants by ion-exchange chromatography.

### SDS-PAGE, western blot and HPLC-SEC

Purified proteins were submitted to SDS-PAGE using SurePAGE (Genscript). Coomassie brilliant blue staining for SDS-PAGE were performed using eStain L1 Protein Staining machine (Genscript). Gels for western blot were transferred onto the nitrocellulose membrane and reacted with COVID-19-convalescent serum (1:500 diluted), followed by washing and reaction with horseradish peroxidase (HRP)-conjugated anti-human pAb. The blots were imaged on FUSION FX7 Spectra multispectral imaging system (VILBER). The size exclusion liquid chromatography (SEC) for purified StriFK protein was performed using a high-performance liquid chromatography system (Waters Alliance HPLC) on a TSKgel G3000PWXL column.

### Surface plasmon resonance (SPR) analysis

The binding affinity of StriFK to recombinant human ACE2 protein was determined by SPR analysis. For the detection, rACE2 (mouse Fc tagged, Sino Biological) proteins were immobilized to a protein G sensorchip (Cytiva) at a level of about 500 response units (RUs) using Biacore 8000 (Cytiva). Serial dilutions of purified StriFK proteins were injected, ranging in concentration from 200 to 0.78 nM. The RU data were fit to a 1:1 binding model using Biacore™ Insight evaluation software.

### Plasmas of convalescent COVID-19 patients

Twelve plasma samples (including 11 convalescent COVID-19 plasmas and one control sample) were used to test their activities to recombinant StriFK protein. The authentic virus neutralizing titers, pseudovirus neutralizing titers, and receptor-blocking titers had been measured and described in our previous study (*44*). Another 268 plasmas collected from recovered COVID-19 patients within eight weeks since illness onset were used to determine the GMT level of neutralizing antibodies (by pseudovirus virus assay) of convalescent COVID-19 patients. All of these convalescent plasmas were collected after the confirmed COVID-19 patients discharged from the First Hospital of Xiamen University. The study was approved by the institutional review board of the School of Public Health in accordance with the Declaration of Helsinki, and written informed consent was obtained.

### Adjuvant characterization

The appearance of FH002C and Al001 was photographed. The morphological characteristics of the two adjuvants were imaged at × 100 using the Motic AE31E inverted microscope (Motic, Xiamen, China), and were observed at a nominal ×120,000 magnification by transmission electron microscopy (JEOL, Tokyo, Japan). The particles’ size and zeta potentials were measured by using LS 13320 Beckman Coulter Laser Particle Analyzer (Beckman Coulter, CA, USA) and NanoBrook Omni zeta potential analyzer (Brookhaven Instruments Corporation, NY, USA), respectively.

### Vaccine formulation

Al001 and FH002C are both suspensions of aluminum-based adjuvants, which were prepared in-house. Al001 is an aqueous solution of amorphous aluminum hydroxyphosphate, much like what was described for adjuvants for HBV, HPV and HEV vaccines (*45-47*), with a molar ratio of P/Al:0.15. The Al content in the formulation is 0.84 mg/mL. The FH002C is a novel functionalized adjuvant, wherein the Al element was partially replaced with zinc, and the amorphous phosphophilic particles (*48*) were loaded with risedronate for additional immunostimulatory activity. The resulting organic-inorganic hybrid adjuvant showed similar whitish suspension morphology in aqueous solution with the widely used A1001 adjuvant (*45*). Before using, the 2x adjuvants were mixed well with resuspending, then each antigen sample (in 150 mM NaCl) was mixed at 2× concentration with an equal volume of the adjuvant to achieve the final desired concentration of antigen and adjuvant in the final formulation. The vaccine formulations were mixed well with several gentle upside-and-downside turns of the vial/tube and stored at 2-8 °C until use. All the vaccine formulations containing Al001 or FH002C were prepared at least 24 hours before immunization. Based on the ELISA results using a sandwich assay with mAbs on both sides, completeness of adsorption was shown to be at least 95% for both formulations based on the antigen content in the supernatant as compared to the overall antigen in the formulation.

### Mouse immunization

For antibody response evaluation, BALB/c mice were maintained in a specific pathogen-free (SPF) environment and immunized with various proteins at 1 μg/dose with Al001 or FH002C through intramuscular injection, following an immunization schedule of one priming dose at Week 0 plus two boosters at Weeks 2 and 4. Serum samples were collected at Week 0, 1, 2, 3, 4, 5, and 6 via retro-orbital bleeding to measure the antibody titers.

For cellular immune response analyses, C57BL/6 mice were immunized 10 μg of StriFK with Al001 or FH002C adjuvants via intramuscular injection on days 0 and 21. Then, splenocytes and lymph node cells were collected on day 28 post-immunization for IFN-γ enzyme-linked immunospot assays (ELISPOT) or intracellular cytokine staining (ICS) measurements. To germinal center response characterization, the T follicular helper cells (Tfh), germinal center B cells (GCB), and plasma cells in LNs from immunized C57BL/6 mice at 1-week after single immunization were analyzed by Flurescence Activated Cell Sorting (FACS).

### Enzyme-linked immunospot assay (ELISPOT)

Briefly, single-cell suspensions from mouse spleen (10^6^ cells per well) or LNs (4×10^5^ cells per well) were seed in precoated ELISPOT plate (Dakewe Biotech). Subsequently, cells were incubated with pooled peptides of SARS-CoV-2 spike (15-mer peptides with 11 amino acids overlap, cover the entire spike protein, Genscript) and cultured for 20 hours. The detecting procedure was conducted according to the manufacturer’s instructions. Spots were counted and analyzed by using CTL-ImmunoSpot® S5 (Cellular Technology Limited). The numbers of IFN-γ secreting cells were calculated by subtracting PBS-stimulated wells from spike peptide pool-stimulated wells.

### Flow cytometry analyses

For GCB cells and plasma cells analyses, lymph node cells were stained with PerCP/cy5.5-conjugated anti-mouse TCRβ (Biolegend, 109228), APC/cy7-conjugated anti-mouse B220 (Biolegend, 103224), PE-conjugated anti-mouse CD95 (BD, 554258), Alexa Fluor 647-conjugated anti-mouse GL-7 (Biolegend, 144606), BV421-conjugated anti-mouse CD138 (Biolegend, 142523), FITC-conjugated anti-mouse IgD (Biolegend, 405704) and LIVE/DEAD™ Fixable Aqua Dead Cell Stain Kit (Invitrogen). For Tfh cells analysis, the following antibodies were used: rat anti-mouse CXCR5 (BD, 551961), biotin-conjugated anti-rat IgG2a (Biolegend, 407504), APC conjugated streptavidin (eBioscience, 17-4317-82), FITC conjugated anti-mouse CD4 (Biolegend, 100406), PE/cy7 conjugated anti-mouse CD8α (Biolegend, 100722), PerCP/cy5.5 conjugated anti-mouse TCRβ (Biolegend, 109228), BV421 conjugated anti-mouse PD-1 (Biolegend, 135221) and LIVE/DEAD™ Fixable Aqua Dead Cell Stain Kit.

For ICS staining, mouse splenocytes (2×10^6^ per test) and lymph node cells (2×10^6^ per test) were stimulated with pooled spike-peptides (1 μg/mL) in a U-bottom plate for an 18-hour incubation in a CO2 incubator. Then, protein transport inhibitors (BD GolgiPlug™, BD Biosciences) were added and incubated with cells for 6 hours. Subsequently, cells were stained with PE/Cy7-conjugated anti-mouse CD4 (Biolegend, 100422), PerCP/cy5.5-conjugated anti-mouse CD8α (Biolegend, 100734), FITC-conjugated anti-mouse CD49b (BD, 553857), and LIVE/DEAD™ Fixable Aqua Dead Cell Stain Kit. Subsequently, cells were fixed and permeated by using Fixation/Permeabilization Solution Kit (BD Biosciences), and further stained with BV421-conjugated anti-mouse IL-4 (Biolegend, 504127) and, PE-conjugated anti-mouse IFN-γ (Biolegend, 505826). Finally, the samples were measured by BD LSRFortessa X-20 Flow Cytometer (BD), and the data were analyzed by FlowJo V10.6.0.

### Anti-RBD, anti-spike, anti-S1, and anti-S2 IgG measurements

Microplates pre-coated with recombinant antigens of RBD, spike ectodomain, S1, or S2, were provided by the Beijing Wantai company. For detections, serial-diluted (2-fold) serum samples (100 μL per well) were added into the wells, and the plates were incubated at 37 °C for 30 minutes, followed by washing with PBST buffer (20 mM PB7.4, 150 mM NaCl and 0.05% Tween 20). Then, HRP-conjugated anti-mouse IgG (for measurements of mouse sera), anti-human IgG (for measurements of monkey sera), or anti-hamster IgG (for measurements of hamster sera) solutions (100 μL per well) were added according to the species of samples. After a further 30-min incubation followed by washing, TMB chromogen solution (100 μL per well) was added into the well. Ten minutes later, the chromogen reaction was stopped by adding 50 μL of 2 M H_2_SO_4_, and the OD450-630 was measured. The IgG titer of each serum was defined as the dilution limit to achieve a positive result (>median+3×SD of ODs of negative controls).

### Measurements for antibody avidities and isotypes in mouse serum

Microplates pre-coated with recombinant spike protein were provided by the Beijing Wantai company. Serial dilutions of serum samples from immunized mice were added into the wells in duplicate microplates. The plates were incubated at 37 °C for 1-hour and followed by a PBST wash cycle (5 times). Then, one of the duplicate microplates was incubated with 100 μL of 4M Urea solution and the other one was incubated with 100 μL of PBS buffer at 37 °C for 20 min. Followed by a wash cycle, HRP-conjugated anti-mouse IgG was added into the wells for incubation of 30 min at 37 °C. Finally, the plates were further washed with PBST for 5 times and followed by chromogen reaction and OD measurements. The binding activity of polyclonal antibodies in serum was defined as the dilution fold required to achieve 50% of the maximal signal (ED_50_). The ratio of ED50(urea-treated) to ED50(PBS-control) was used to measure the avidity of serum polyclonal antibodies to StriFK.

Serum IgG isotype-specific antibody titers were determined by a modified ELISA method. For the assay, recombinant spike coated microplates were used. The serum samples were serially diluted and reacted with plates following a similar procedure as above mentioned in IgG titer measurements. After a 1-hour incubation, the plates were washed for 5 times and subsequently incubated with HPR-conjugated goat anti-mouse IgG1, IgG2a, or IgG2b (AbD Serotec, Oxford, UK) at 37 °C for 30-minutes. Then, the plates were washed for 5-time and followed by chromogen reaction and OD measurements. The anti-spike titers of IgG1, IgG2a, and IgG2b were determined as the dilution limit to achieve a positive result.

### Pseudovirus neutralizing antibody titer measurement

Vesicular stomatitis virus (VSV) based SARS-CoV-2 pseudotyping-particles (VSVpp) were produced according to our previous study (*49*). The BHK21-hACE2 cells were used to perform the pseudovirus neutralization assay. Briefly, cells were pre-seeded in 96-well plates. Serially-diluted (3-fold gradient) samples were incubated with VSVpp inoculum for 1 hour. Next, the mixture was added into seeded BHK21-hACE2 cells. After a further incubation at 37 °C containing 5% CO2 for 12 hours, fluorescence images were captured by Opera Phenix or Operetta CLS High-Content Analysis System (PerkinElmer). The counts of GFP-positive cells of each well were analyzed by the Columbus System (PerkinElmer). The neutralization titer of each sample was expressed as the maximum dilution fold (ID50) required to achieve infection inhibition by 50% (50% reduction of GFP-positive cell numbers compared to controls).

### Receptor-blocking antibody titer measurement

A cell-based spike function-blocking test (CSBT) was used to characterize the receptor-blocking antibody titer in animal serum following the previously described procedure (*44*). Briefly, the hACE2-mRuby3 293T (293T-ACE2iRb3) cells were seeded in poly-D-lysine pretreated CellCarrier-96 Black plate at 2×10^4^ cells per well overnight. Hamster or Cynomolgus monkey sera were diluted in a 2-fold serial dilution with DMEM medium containing 10% FBS. 11 μL Gamillus-fused SARS-CoV-2 spike trimer (STG) probe mixed with 44 μL of the diluted sera to a final concentration of 2.5 nM. After removing half of the medium from the pre-seeded 293T-ACE2iRb3 cells, pipette 50 μl of the mixture into the wells and incubate 1 hour at 37 °C containing 5% CO2. Subsequently, cell images were captured by Opera Phenix High-Content Analysis System in confocal mode. Quantitative image analyses were performed by the Columbus System (PerkinElmer) following the previously described algorithm (*44*). The CSBT activities (receptor-blocking antibody titers) of immunized sera were displayed as ID50, calculated by 4PL curve-fitting in GraphPad Prism.

### Hamster vaccination

Fifty Syrian golden hamsters (male:female=1:1) were used for vaccine immunization and the SARS-CoV-2 challenge *in vivo*. For 2-dose vaccination, the vaccines containing 10 μg antigen (RBD or StriFK) formulated with FH002C adjuvant were used to immunize hamsters (n=6 per group) at Weeks 0 and 2. Another group was vaccinated with FH002C adjuvant without antigen as a sham control (n=6). For 3-dose vaccination, groups of hamsters (n=16 per group) received 10 μg StriFK formulated with Al001 or FH002C adjuvants at Weeks 0, 2, and 6. All hamsters received 200 μL of vaccine injection per dose via the intramuscular route. Serum samples were collected weekly for antibody analyses.

### Cynomolgus monkey vaccination

Eight Cynomolgus monkeys were allocated randomly into four groups (one female and one male per group). Groups of cynomolgus were injected with 20 μg StriFK with Al001 or FH002C, or 20 μg RBD with FH002C adjuvant per dose via the intramuscular route for 3 doses. Another group received FH002C adjuvant-only as a sham control. All cynomolgus were injected 0.5 mL of vaccine or adjuvant-only at Weeks 0, 2, and 6. Serum samples were collected weekly for biochemical and antibody analyses, including measurement of anti-spike IgG, pseudovirus neutralizing antibody, and receptor-blocking antibody titers during the Week 1 to 10. Cynomolgus monkey immunogenicity experiment was conducted at JOINN Laboratories, Inc (Beijing).

### SARS-CoV-2 virus challenges in hamsters

Both intranasal challenge and direct contact challenge of SARS-CoV-2 were used in the study. For the former, hamsters were inoculated with 1×10^4^ PFU of the SARS-CoV-2 virus through the intranasal route under anesthesia. For the latter, virus-carried hamsters (donors) were pre-infected with inoculation of 1×10^4^ PFU of the SARS-CoV-2 virus through the intranasal route. One day later, every four donors were transferred to a new cage and were co-housed with four vaccinated or unvaccinated control animals. The hamsters were fed with a daily food amount of 7 g per 100 g of body weight. The weight changes and typical symptoms (piloerection, hunched back, and abdominal respiration) in hamsters were recorded daily since the infection or contact. Hamsters may be sacrificed for tissue pathological and virological analyses on days 3, 5, or 7 after the virus challenge. The virus challenging studies were performed in the anmial biosafety level 3 (ABSL-3) facility.

### SARS-CoV-2 RNA quantification

Viral RNA levels in lung, trachea, and nasal turbinate from challenged hamsters were detected by quantitative RT-PCR. Hamster tissue samples were homogenized by TissueLyser II (Qiagen, Hilden, Germany) in 1mL PBS. The RNA was extracted using the QIAamp Viral RNA mini kit (Qiagen) according to the manufacturer’s instruction. Subsequently, viral RNA quantification was conducted using SARS-CoV-2 RT-PCR Kit (Wantai, Beijing, China).

### SARS-CoV-2 plaque assay

The plaque assays were performed in the biosafety level 3 (BSL-3) facility. Briefly, Vero cells were seeded in a 6-well plate at 10^5^ cells per well. After overnight culture, the medium was removed, and the cells were washed twice with PBS. The supernatants of tissue lysates were serially 10-fold diluted with DMEM and were incubated with the cells. Cell plates were then incubated for 1 hour at 37 °C containing 5% CO_2_. Subsequently, the tissue lysate contained medium was refreshed with a mixture of DMEM containing 1% agar. After agar solidification, the cell plates were inverted and incubated at 37 °C containing 5% CO_2_ for 3 days. Subsequently, cells were fixed with 2 mL of 4% formaldehyde for 2 hours at room temperature. The cells were further stained with 0.1% crystal violet for 20 min. The viral load was calculated based on the count of plaques and the corresponding dilution factor.

### Histopathology

The lung tissues from challenged hamsters were fixed with 10% formalin for 48 hours, then embedded in paraffin and sectioned. Next, the fixed lung sections were subjected to hematoxylin and eosin (H&E) staining. Immunohistochemical staining was performed by using the mouse monoclonal anti-SARS-CoV-2 N proteins antibody. Whole-slide images of the lung sections were captured by EVOS M7000 Images System (ThermoFisher).

### Statistical analysis

The Student’s t-test or Mann-Whitney U test was used for comparison of continuous variables between groups. The two-way ANOVA test was used to analyze the time-serial observations for the independent sample. Pearson test and linear regression model were used for univariate correlation analysis. Statistical significance was considered to be significant for 2-tailed p values <0.05. Statistical analyses were conducted in GraphPad Prism 8 software.

## Acknowledgements

This study was supported by National Natural Science Foundation of China (81991491, 31730029, U1905205, 81702006, 81902057 and 81871316), National Key Plan for Scientific Research and Development of China (2016YFD0500302; 2017YFE0190800), funding for Guangdong-Hongkong-Macau Joint Laboratory (2019B121205009).

## Author Contributions

Project conceptualization: J.Z., T.Y.Z., H.C.Z., Q.J.Z., Q.Y., Y.G., N.S.X.; Constructs design, protein production and characterization: S.J.W, M.W., Y.T.W., Y.L.Z., Z.L.L., Q.B.Z., Q.Y.; Adjuvant design, production and characterization: X.F.H., M.F.N., Z.G.Z., M.X.Y., Q.J.Z.; Animal studies: Y.T.W., L.Z.Y., R.R.C., J.M., J.J.X., L.Q.C., M.Z., Y.S., M.P.C., J.P.H. H.C.Z.; Antibody response analysis: Y.L.Z., H.L.X., X.F.H, Y.T.W., Z.K.W, Y.D.H., D.B.C.; Cellular immune response analysis: Y.T.W., R.Y.Q., L.Z., T.Y.Z.; Manuscript writing: Y.T.W., X.F.H., T.Y.Z., H.C.Z., Q.J.Z., Q.Y.; Critical revision: S.W.L., Y.X.C., T.C., J.Z., Y.G., N.S.X.; Approved the final version of the manuscript: J.Z., T.Y.Z., H.C.Z., Q.J.Z., Q.Y., Y.G., N.S.X.

---

**Figure S1.**
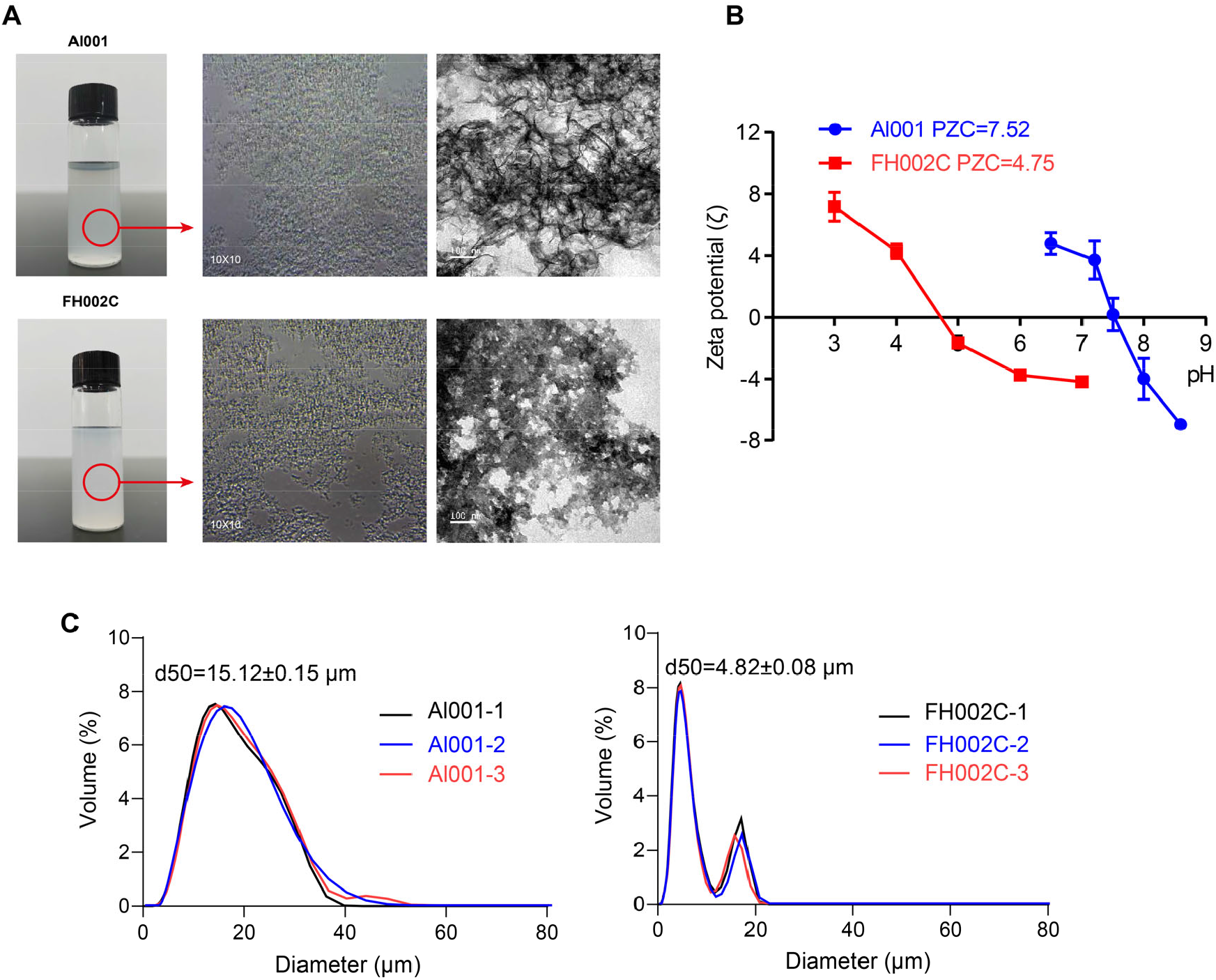
Characterizations of the Al001 and FH002C adjuvants. (A) The appearance and morphology of Al001 (upper panel) and FH002C (lower panel) observed by cell-phone camera (left), inverted microscope (middle), and transmission electron microscopy (right). The potential of zero charge (PZC) of Al001 and FH002C. Zeta potential of Al001 and FH002C were measured in different pH using NanoBrook Omni zeta potential analyzer (Brookhaven Instruments Corporation, NY), and the PZC values were calculated by the instrument software. Each Zeta potential value reported is the average of three replicates. Size distributions of Al001 (left panel) and FH002C (right panel). The particle size distribution of adjuvants was measured by using LS 13320 Beckman Coulter Laser Particle Analyzer (Fullerton, CA). The median particle diameter d_50_ (equivalent diameter when the cumulative distribution is 50%) was measured in three independent replicates, mean±SEM.

**Figure S2.**
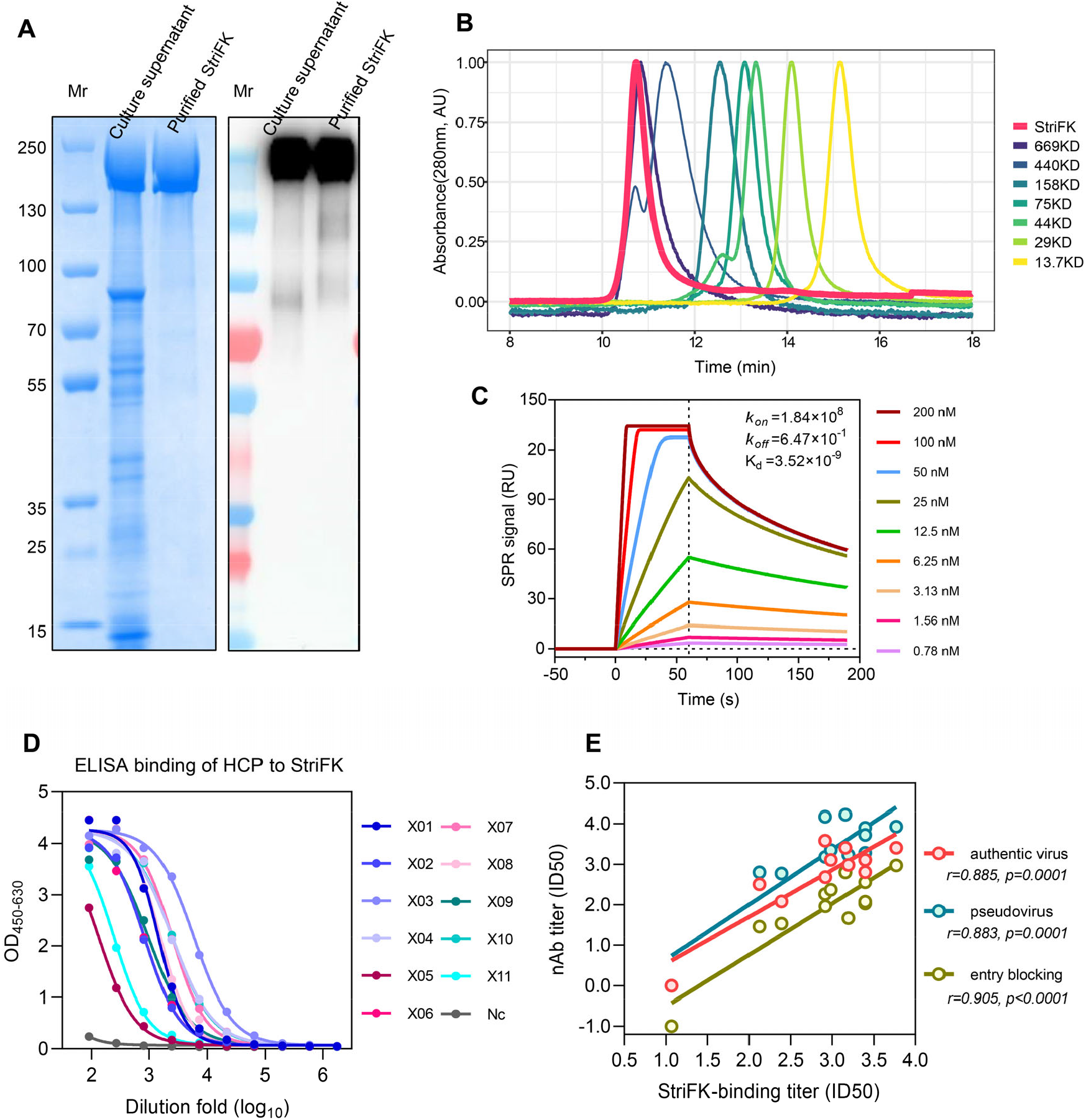
Characterizations of recombinant StriFK protein. (A) Stable CHO cell pool produced recombinant StriFK at a high level. SDS-PAGE and western blot analyses for culture supernatants (unconcentrated) and purified protein derived from an StriFK-integrated stable CHO pool. (B) SEC chromatograms of StriFK protein with protein standards on a TSK-G3000 column. (C) SPR sensorgrams showing the binding kinetics for StriFK with immobilized human ACE2. Colored lines represented a global fit of the data using a 1:1 binding model. (D) ELISA binding activities of human COVID19-convalescent plasmas to StriFK protein. Twelve representative samples (11 convalescent COVID-19 plasmas and 1 control sample) were used for the tests. (E) Correlation analyses between the StriFK-binding titer and the pseudovirus neutralizing antibody titer (colored in blue), the authentic SARS-CoV-2 virus-neutralizing antibody titer (colored in red), and the receptor-blocking antibody titer (colored in yellow) among human plasmas.

**Figure S3.**
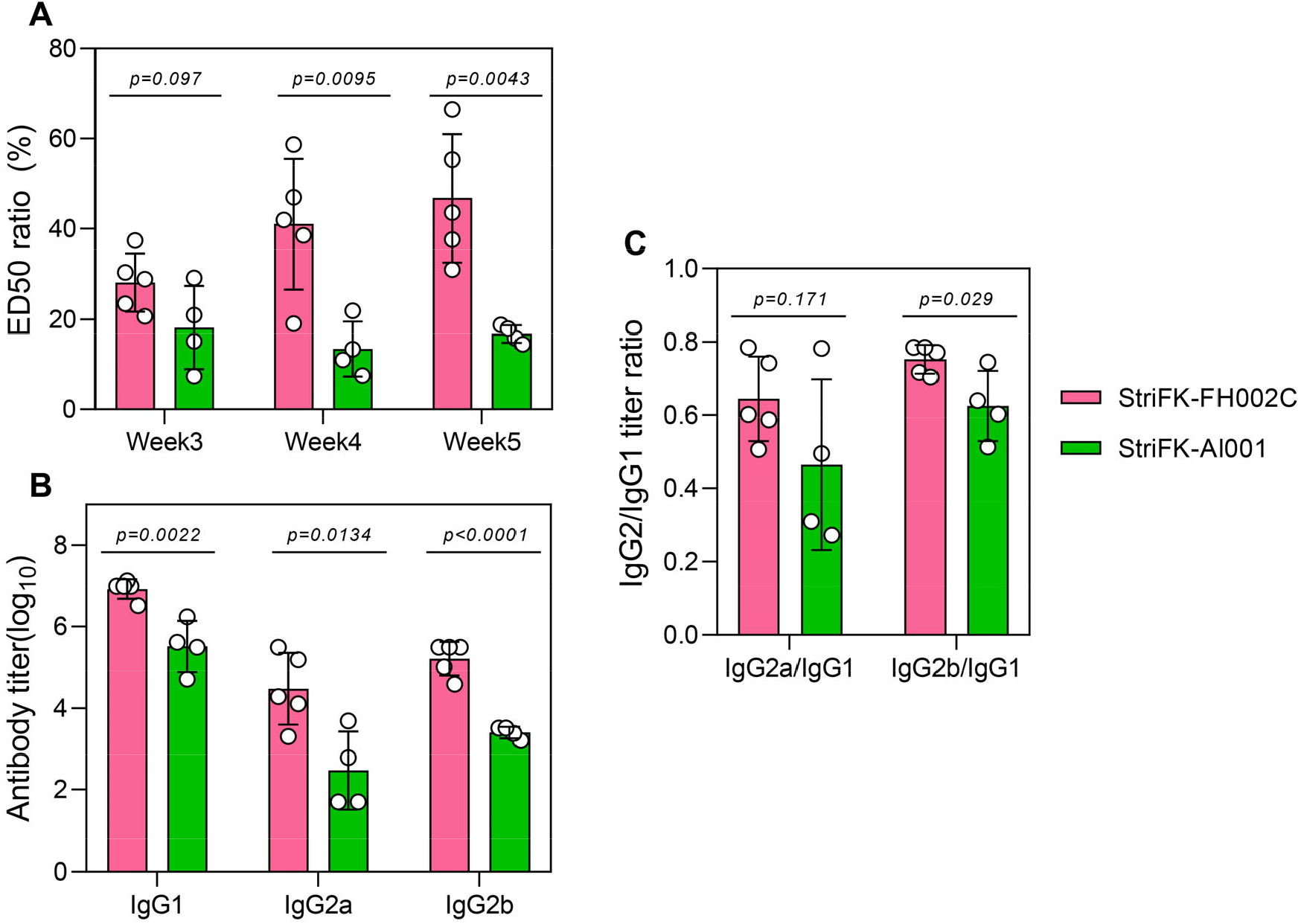
The IgG avidity and IgG subclass of spike-specific antibodies induced by StriFK-FH002C and StriFK-Al001 in mice. BALB/c mice were immunized at Weeks 0 and 2 with 1 μg/dose of StriFK-FH002C (n=5) or StriFK-Al001 (n=4). (A) Serum IgG avidities of immunized mice at Weeks 3, 4, and 5. The ratio of ED_50_ (urea-treated) to ED_50_ (untreated control) was used to measure the avidity of the serum polyclonal antibody to StriFK. (B) Serum anti-spike binding titers of IgG1, IgG2a, and IgG2b for immunized mice. Sera were collected 2 weeks post the 2^nd^ dose administration. (C) Antibody titer ratios of IgG2a to IgG1 and IgG2b to IgG1 were calculated. Unpaired *t*-test was used for statistical comparison. Related to Figure 1C.

**Figure S4.**
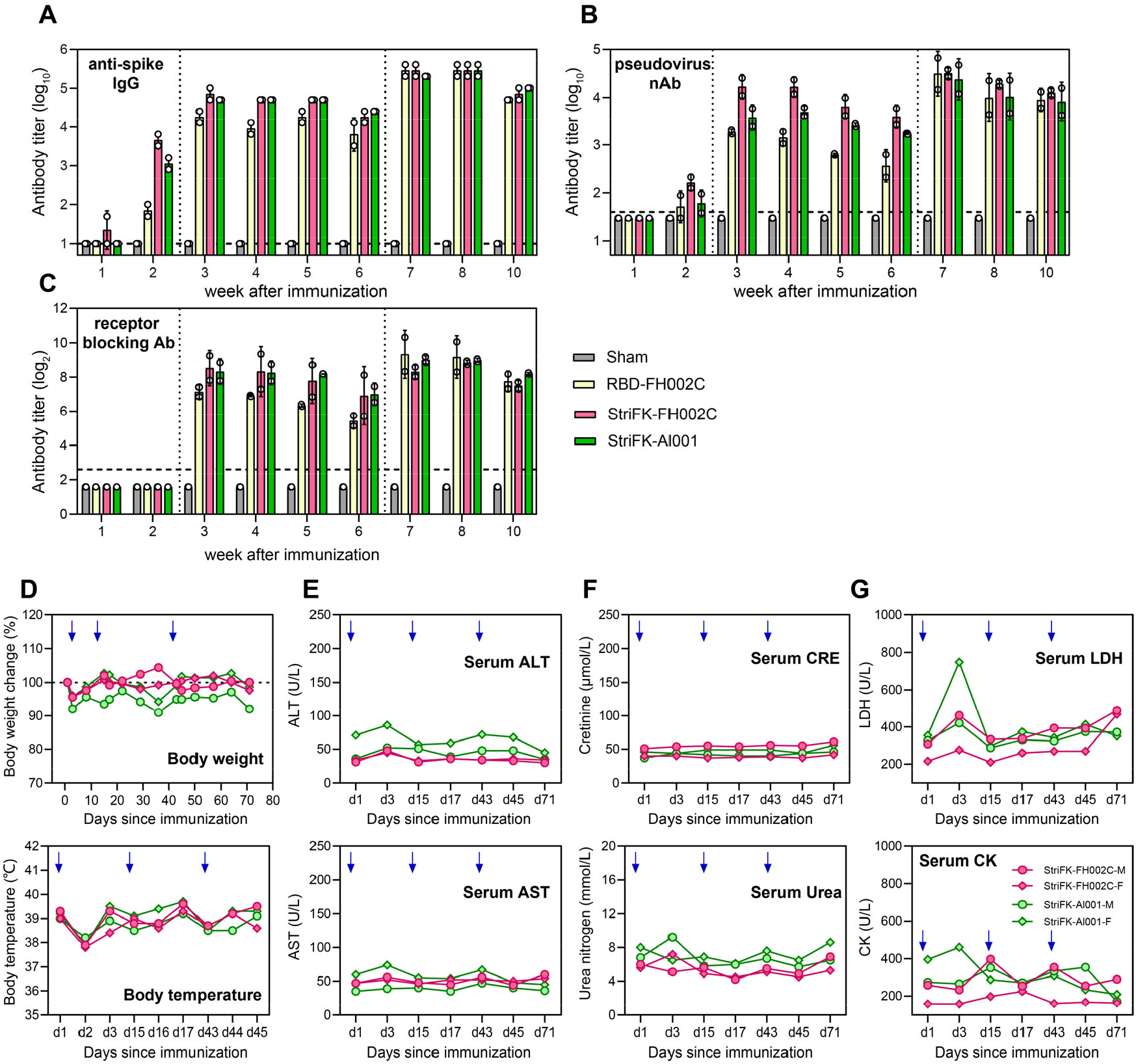
Immunogenicity and safety profiles of vaccine candidates in non-human primates. For each vaccine, two cynomolgus macaques were immunized for 3 shots of vaccine candidates (20 μg/dose) at Weeks 0, 2, and 6. Serum samples were collected every week from Week 1 to 10 after initial immunization, and the anti-spike (A), neutralizing antibody (B), and the receptor-blocking antibody titers (C) were measured. (D) The body weight (upper panel) and temperature (lower panel) of cynomolgus macaques following StriFK-FH002C and StriFK-Al001 immunizations. Serum biomarkers of liver function (ALT and AST, panel E), renal function (Creatinine and Urea nitrogen, panel F), and cardiac enzymes (LDH and CK, panel G) of cynomolgus macaques following StriFK-FH002C and StriFK-Al001 immunizations.

**Figure S5.**
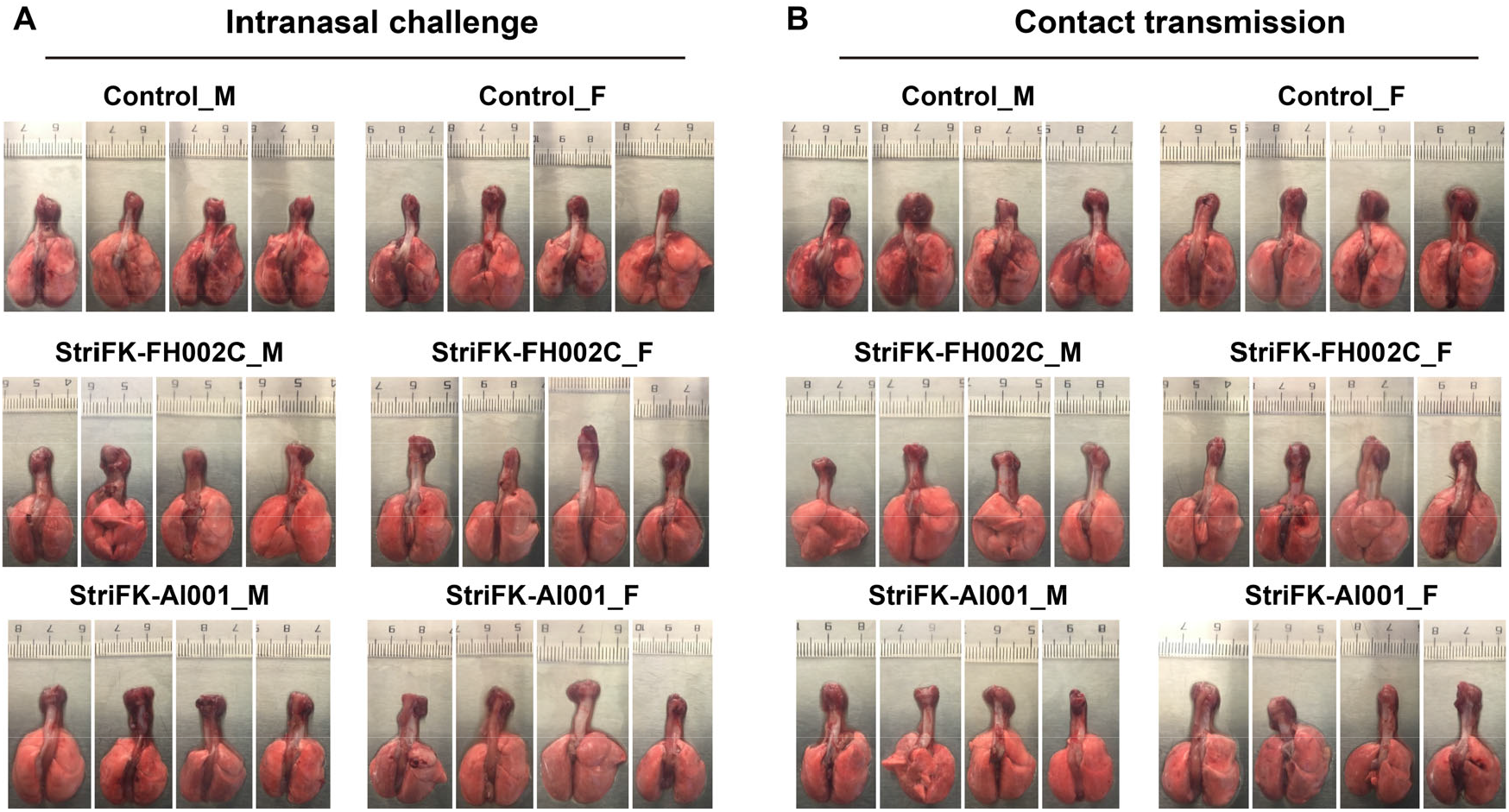
Gross observations (upper panel) and infectious viral titers (lower panel) of lung tissues collected from vaccinated and control hamsters at 5dpi after SARS-CoV-2 challenges. (A) Hamsters were challenged via intranasal inoculation of SARS-CoV-2. Related to Figure 5. (B) Hamsters were exposed to SARS-CoV-2 via direct contact with pre-infected donor hamsters. Related to Figure 6.

**Figure S6.**
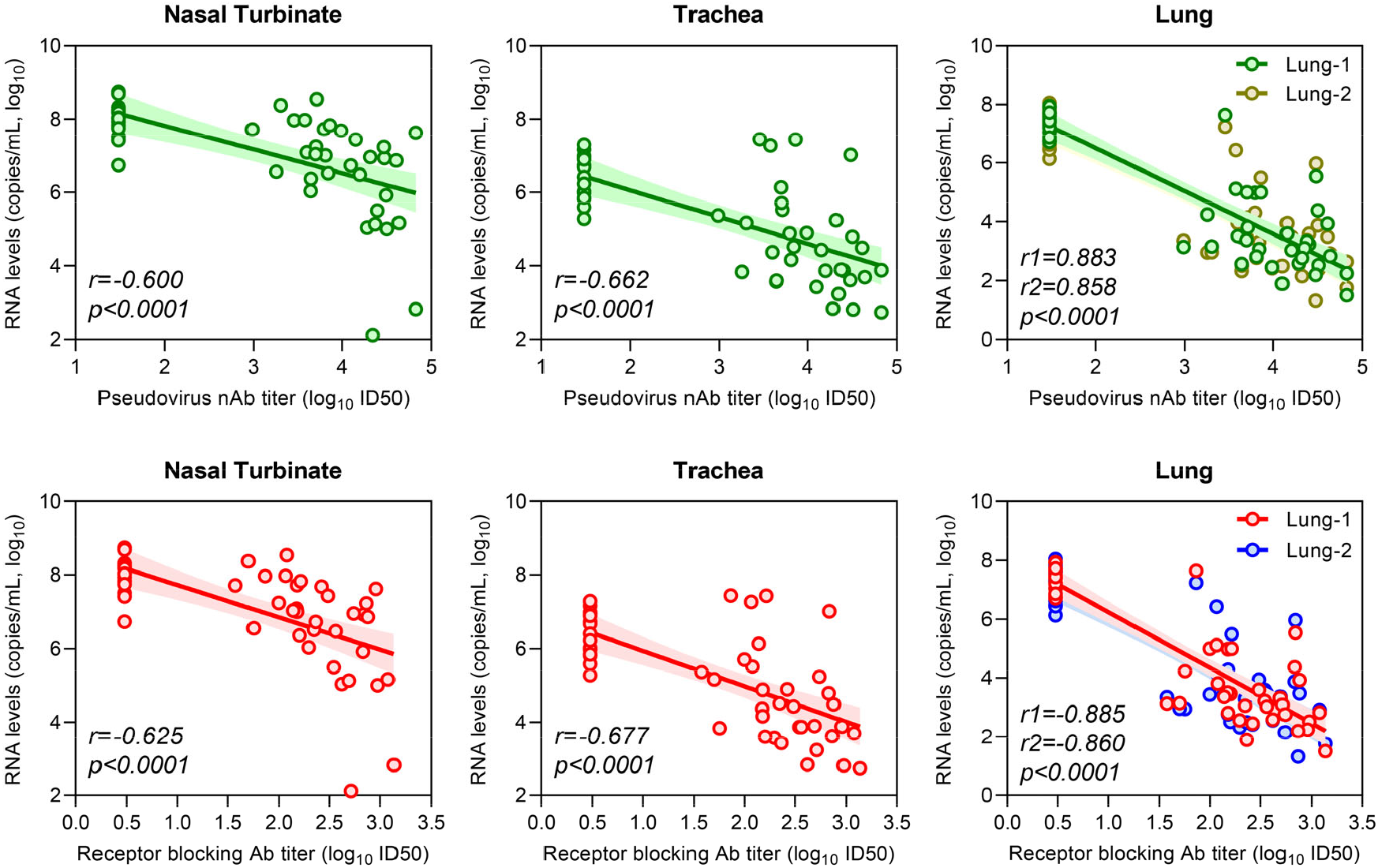
Correlations between the tissue viral RNA levels and serum pseudovirus nAb (upper panels) or receptor-blocking antibody (lower panels) titers among SARS-CoV-2 challenged hamsters. The antibody titers of these hamsters at Week 7 (3 days before virus challenging study) since initial immunization were used for analysis. Tissue viral RNA levels at 5 dpi (as shown in Figure 5C and Figure 6C) were used. All vaccinated hamsters and unvaccinated controls involved in experiments related to Figures 5 and 6 were included and pooled for analysis. Lung-1 and Lung-2 indicated lung tissues collected from tissues near and away from the pulmonary hilum, respectively.

